# Molecular cartography uncovers evolutionary and microenvironmental dynamics in sporadic colorectal tumors

**DOI:** 10.1101/2023.03.09.530832

**Authors:** Cody N. Heiser, Alan J. Simmons, Frank Revetta, Eliot T. McKinley, Marisol A. Ramirez-Solano, Jiawei Wang, Justin Shao, Gregory D. Ayers, Yu Wang, Sarah E. Glass, Harsimran Kaur, Andrea Rolong, Bob Chen, Paige N. Vega, Julia L. Drewes, Nabil Saleh, Simon Vandekar, Angela L. Jones, M. Kay Washington, Joseph T. Roland, Cynthia L. Sears, Qi Liu, Martha J. Shrubsole, Robert J. Coffey, Ken S. Lau

## Abstract

Colorectal cancer exhibits dynamic cellular and genetic heterogeneity during progression from precursor lesions toward malignancy. Leveraging spatial molecular information to construct a phylogeographic map of tumor evolution can reveal individualized growth trajectories with diagnostic and therapeutic potential. Integrative analysis of spatial multi-omic data from 31 colorectal specimens revealed simultaneous microenvironmental and clonal alterations as a function of progression. Copy number variation served to re-stratify microsatellite stable and unstable tumors into chromosomally unstable (CIN+) and hypermutated (HM) classes. Phylogeographical maps classified tumors by their evolutionary dynamics, and clonal regions were placed along a global pseudotemporal progression trajectory. Cell-state discovery from a single-cell cohort revealed recurring epithelial gene signatures and infiltrating immune states in spatial transcriptomics. Charting these states along progression pseudotime, we observed a transition to immune exclusion in CIN+ tumors as characterized by a novel gene expression signature comprised of *DDR1, TGFBI, PAK4,* and *DPEP1*. We demonstrated how these genes and their protein products are key regulators of extracellular matrix components, are associated with lower cytotoxic immune infiltration, and show prognostic value in external cohorts. Through high-dimensional data integration, this atlas provides insights into co-evolution of tumors and their microenvironments, serving as a resource for stratification and targeted treatment of CRC.

## Introduction

The genetic model of colorectal cancer (CRC) progression defines a sequence of cumulative mutational burden that drives dysplasia and malignancy in human colonic epithelium^1^. This conventional adenoma-carcinoma trajectory involves alteration of driver genes *APC, KRAS,* and *TP53,* resulting in chromosomal instability (CIN)^2^. Alternatively, a subset of CRCs which arise from the so-called serrated pathway, are more likely to be BRAF-driven and microsatellite unstable (MSI-H) due to hypermethylation of *MLH1* and other mismatch repair (MMR) genes, causing hypermutation^3,4^. Ensuing decades of investigation have yielded additional CRC subtyping that elucidates alternative pathways to invasion and metastasis, as well as characterization of pre-malignant lesions and their clinical prognoses^5–7^. More recently, the advent of single-cell and spatial molecular assays has uncovered various degrees of intratumoral heterogeneity at high resolution, suggesting that previously proposed linear tumor progression along the conventional or serrated pathway cannot fully explain the evolutionary dynamics of the second leading cause of cancer-related mortality world-wide^8–11^. Moreover, layered molecular information from spatially resolved assays can be used to build models relating gene and protein expression, or cell “state”, to clonal identity across tumor regions^12–15^. Exploration of tumor phylogeography in this way allows for deeper profiling of evolutionary relationships while accounting for regional heterogeneity^16^. Beyond, and perhaps fundamental to, genetic evolution of malignant cells themselves lies additional complexity introduced by interactions between tumor epithelium and infiltrating immune cells, which provide immunogenic selection pressure that profoundly impacts tumor evolution and prognosis^17–19^. Indeed, characterization of the immune compartment of solid tumors has been shown to be a better predictor of patient prognosis than traditional pathological staging, and tumor immunophenotype is a valuable measure in forecasting response to immune checkpoint inhibition (ICI) in several cancers ^20,21^. Furthermore, distinct pathways from initiation to malignancy confound these tumor characteristics, as serrated lesions and MSI-H CRCs are more immunogenic than their conventional adenoma and microsatellite stable (MSS) counterparts on average^4,9^. Importantly, several types of advanced solid tumors have been shown to exhibit immune exclusion or evasion, by mechanisms both intrinsic to cancer cells and observed microenvironmentally, which shortens overall patient survival and confers ICI resistance^22–29^. In CRC, an observed suppression of cytotoxic immunity that trends with an increased stem cell signature in late-stage carcinoma raises a critical question surrounding immune exclusion and the potential connection to tumor progression^9^.

In order to map the co-evolution of colorectal tumor cells and their microenvironments, we leveraged spatial multi-omics to build a phylogeographical atlas of CRC progression from pre-cancer to adenocarcinoma. We have generated a novel, spatially resolved dataset from a heterogeneous set of sporadic colorectal tumors, wherein distinct tumor regions represent snapshots of cancer evolution. Multiregional mutational profiling, untargeted spatial transcriptomics (ST), and multiplexed protein imaging offer high-dimensional paired measurements of layered molecular information while maintaining tissue contexts^30–32^.

Combining spatial data with single-cell RNA sequencing (scRNA-seq) to enumerate consensus cell states, we projected tumor programs and microenvironmental features onto a generalized progression pseudotime (PPT) derived from regional copy number variants (CNVs) and somatic mutational profiles amongst a cohort of tumors. These efforts enabled discovery of multiple pathways that are distinctly altered during the progression of chromosomally unstable (CIN+) and hypermutated (HM) CRCs. This study presents a patient-centric roadmap of CRC evolution and progression arising from integrative, atlas-wide analyses across a unique and heterogeneous set of human CRC specimens.

## Results

### Spatial atlas queries layers of molecular heterogeneity in sporadic colorectal tumors

To model CRC progression through spatial heterogeneity, we selected human colonic specimens with regional morphologies representing transitions between tumor progression stages. Samples with concurrent pre-malignant, malignant, and invasive regions were identified by a pathologist (Methods: Sample procurement). A diversity of tumor stages, grades, and locations was represented (Figure 1A-B; Table S1) with fairly equal selection of MSS (*n* = 12) and MSI-H (*n* = 10) CRCs and pre-cancerous polyp subtypes (*n* = 8; 4 SSL/HP and 4 TA/TVA). Along with a normal colon sample as control, these specimens provide a representative spectrum of disease states along the two major pathways to malignancy in the colon (Figure 1A-B; Table S1).

**Figure 1.**
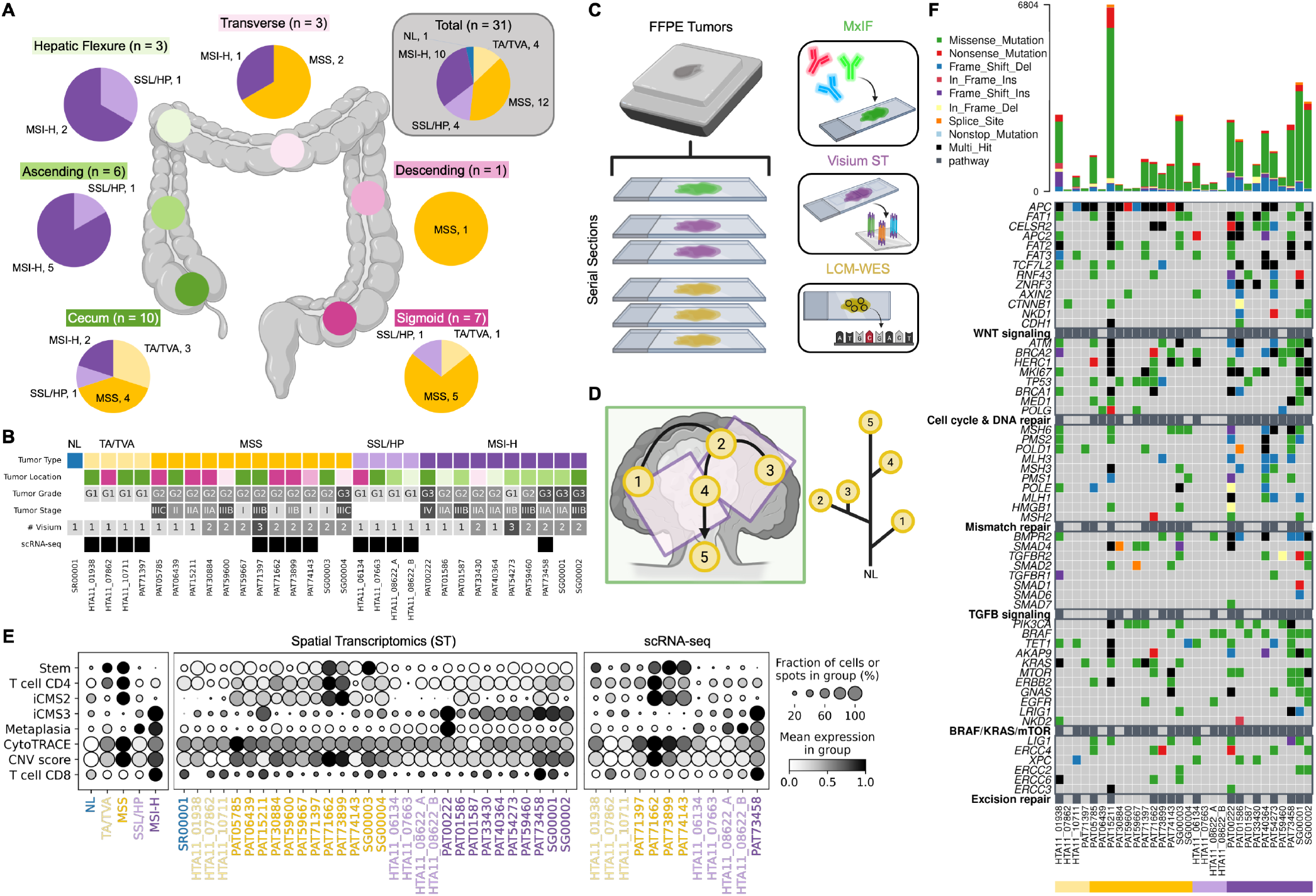
Spatial atlas queries layers of molecular heterogeneity in sporadic colorectal tumors. (A) Diagram detailing colorectal specimens chosen for atlas experiments. (B) Patient-level information from A in table format. Large CRCs (> 7mm diameter; 12 MSS, 10 MSI-H) have at least 1 spatial transcriptomics (ST) replicate tiling the tissue. TA/TVAs (4), HP/SSLs (4), and NL (1) have 1 ST replicate each. All specimens have whole-slide MxIF imaging and bulk or multiregional WES. (C) Experimental design consisting of layered spatial molecular assays from serial sections of FFPE tissue blocks. (D) Diagram of phylogeographical cartography from multiregional sequencing (LCM-WES) data, layered with ST and MxIF images. Black arrows represent progression pseudotime (PPT) inferred from the phylogenetic relationships between ROIs. (E) Summary of gene signature scores by tumor type (left), ST patient (middle), and matched scRNA-seq patient (right). Patient ID colors represent tumor type (MMR status). Mean signature expression scaled across groups. In this dotplot and hereafter, size of dots represents expression frequency, while shade represents intensity. (F) Somatic mutations detected in LCM-WES samples, summarized by patient and grouped by biological pathway. Top barplot represents overall TMB breakdown by mutation class per patient.

For each of these 31 formalin-fixed, paraffin-embedded (FFPE) specimens, we collected serial tissue sections for parallel processing by molecular profiling assays. Multiplex immunofluorescence (MxIF) analysis with a 33-marker panel provided whole-slide protein expression at subcellular resolution^33^ (Figure 1C). We cut serial sections into one or more capture areas of 10X Genomics Visium spatial transcriptomics (ST) slides^34,35^. Large tumors (> 7mm diameter) were trimmed to regions of interest (ROIs) targeting dysplastic epithelium and minimizing stromal areas. H&E images were collected to align gene expression with tissue morphology and provide fiducial markers for spatial registration to MxIF. We employed laser capture microdissection followed by whole-exome sequencing (LCM-WES) on additional serial sections to genetically profile ~2mm ROIs in spatially distinct regions of individual tumors (Figure 1C). Dissociative single-cell RNA sequencing (scRNA-seq) for a subset of specimens provided a reference for expected cellular composition^36,9^.

The data procured from multi-modal analysis of adjacent sections present trade-offs across large ranges of spatial resolution and molecular specificity: MxIF offers subcellular (< 0.5 μm) imaging of dozens of molecular features, ST captures up to 19,000 mRNA transcripts at 100 μm spatial resolution, and LCM-WES profiles somatic mutations for large (1-2mm) regions of tissue. We used image processing software to register genetic, transcriptomic, and proteomic organizational layers, modeling these molecular mixtures at varying spatial scales (Methods: Spatial registration). We use these spatial features to infer relationships of tumor progression and evolution (Figure 1D), and thus provide a “scaffold” for phylogenetic cartography and subsequent modeling of gene, protein, and cellular features^16^.

RNA expression data from 48 ST samples and a subset of 13 matched scRNA-seq samples largely exhibited expected gene expression trends across tumor class and grade when analyzed at the bulk level. Conventional TA/TVA polyps and MSS CRCs were enriched for stemness, intrinsic consensus molecular subtype 2 (iCMS2) epithelial signature, and a CD4+ T lymphocyte-dominant immune response, while SSL/HP and MSI-H CRCs were comprised of metaplastic and iCMS3 signatures accompanied by cytotoxic (CD8+ T cell) immunity^9,10^ (Figure 1E; Table S2). Additionally, aggregating multiregional exome sequencing into bulk analyses per patient revealed *APC, KRAS,* and *TP53* mutations in TA/TVA/MSS tumors, consistent with the conventional adenoma-carcinoma sequence, and *BRAF* variants in SSL/HP/MSI-H samples (Figure 1F). Similar to previous observations, MSI-H CRCs exhibit hypermutation that is absent in their SSL/HP counterparts, delineating that this transition occurs after serrated pre-malignancy^9^. Moreover, a small subset of conventional pathway tumors (1 TVA - HTA11_01938 and 1 MSS - PAT15211) can be identified as hypermutated (HM), consistent with previous observations, likely due to deficiency in proofreading polymerases^37^ (*POLE, POLD1*; Figure 1F).

### CNV inference establishes spatially resolved tumor clones and their phylogenetic relationships

A hallmark of the conventional adenoma-carcinoma sequence that gives rise to MSS CRC is chromosomal instability (CIN) which results in somatic gains, losses, and rearrangements of large segments of DNA heritable to cellular progeny^4,38^. Thus, cumulative increases in CNVs can be used to order tumor progression events amongst tumor regions if measured spatially^39^.

We inferred CNVs from ST data, quantifying levels of CIN across the atlas with support from orthogonal measurements including scRNA-seq, WES, and WGS data^40^. Dimension reduction and embedding of inferred CNV profiles from epithelial ST microwells yields tumor-specific clustering indicative of unique somatic CNVs (Figure 2A). A subset of patients with matched scRNA-seq (*n* = 11) exhibited similar CNV profiles inferred from both scRNA-seq and ST, with exception of CIN-low pre-malignant tumors whose inferred copy number changes consisted of mostly background noise (Figure S2A-D). To quantify this copy number validation between gene expression modalities, we calculated pairwise cosine similarities between inferred CNV profiles of all cells (scRNA-seq) and microwells (ST) assigned to major clones in each tumor (Methods: CNV inference from ST and scRNA-seq). The resulting distributions of these CNV similarities describes the confidence of inferred somatic copy number calls, as chromosomally unstable (CIN+) tumors have cosine similarity distributions approaching 1.0, while CIN-specimens have cosine similarities centered around 0.0 or less (Figure S2B-D). These results derive from the fact that CIN-specimens have so few somatic CNVs that the attempted inference of such rearrangements by expression yields low-confidence, noisy results in both modalities (Figure S2D).

**Figure 2.**
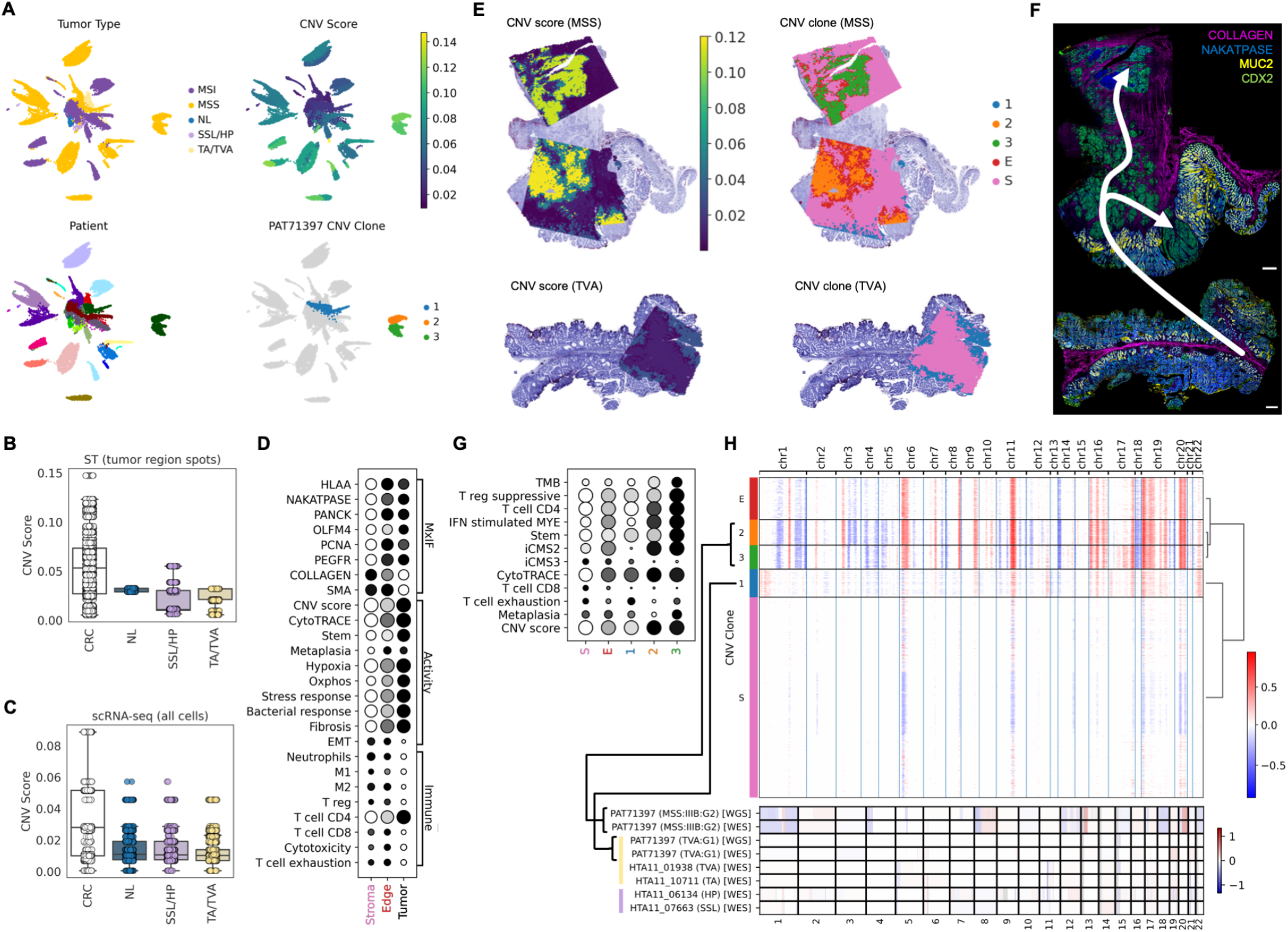
CNV inference establishes spatially resolved tumor clones and their phylogenetic relationships. (A) UMAP embeddings generated from inferred CNV profiles of all ST samples colored by tumor type, CNV score, Patient, and PAT71397 CNV clone to accompany panels E-H. Individual points represent ST microwells, which were subsetted to major clone regions prior to embedding. (B) Boxplots of CNV scores for all ST microwells in major CNV clone regions across atlas, grouped by sample type. CRC *n* = 48,439; NL *n* = 1,067; SSL/HP *n* = 1,735; TA/TVA *n* =1,951. (C) Boxplots of CNV scores for epithelial cells from Chen, *et al.* cohort, grouped by sample type. CRC *n* = 11,982; NL *n* = 31,917; SSL/HP *n* = 11,896; TA/TVA *n* = 21,275. (D) Summary of MxIF intensities, cell activity, and immune gene signatures by major tissue domains determined through CNV inference. (E) CNV scores (left) and tumor clone regions (right) for PAT71397. (F) MxIF with inferred progression trajectory for PAT71397. Scale bars 500 μm. (G) Summary of TMB, CNV score, and gene signatures for CNV clone regions of PAT71397. (H) Heatmap of inferred CNVs for PAT71397 ST, corresponding to E-G (top), as well as CNVs measured by WGS and WES for PAT71397 blocks and additional selected pre-malignant tumors (bottom). Brackets connect WES and WGS from PAT71397 malignant (MSS) and benign (TVA) blocks to dominant CNV clones in respective ST to show similarity of measured and inferred CNV profiles.

To further validate these results using direct measurements of genomic alterations, we performed whole-genome (WGS) and/or whole-exome (WES) sequencing on a subset of tumors from this atlas, as well as additional pre-malignant lesions and CRCs from a larger cohort^9^. CNVs called in these data confirmed somatic copy number alteration patterns inferred by ST and scRNA-seq in overlapping samples (Figure S2E; Methods: CNV calling from WES and WGS).

Several studies have demonstrated that APC dysfunction causes CIN due to the protein’s interaction with microtubules in the spindle and contractile ring during cytokinesis^41,42^. Thus, CIN has been thought to arise as an early event in tumorigenesis, potentially when APC function is lost during initial adenoma formation, as implicated in mouse models^43^. However, human studies using limited numbers of specimens do not provide a clear confirmation^44,45^. To address the relationship between APC and CIN on a broader basis in humans, we summarized CNV scores across our ST atlas and a large cohort of scRNA-seq derived from CRCs and pre-cancers^9^ (*n* = 85), demonstrating that conventional adenomas (TA/TVA) harboring *APC* mutations exhibited low CIN comparable to serrated polyps (SSL/HP) and baseline normal epithelium (Figure 2B-C). We performed WGS (*n* = 35) and WES (*n* = 18) on a selected subset of tumors and similarly calculated total CNV scores from these data, which validated the lack of CIN in TA/TVAs inferred from gene expression (Figure 2H; Figure S2E-F; Methods: CNV calling from WES and WGS).

Taken together, these data suggest that the onset of CIN occurs later in carcinoma development than previously assumed, and that MSS tumors are more likely to become CIN+ than MSI-H carcinomas are^2,46,44,47,48^. We do, however, observe some MSI-H tumors that gain CIN, likely coincident with an observed transformation to a stem-like, iCMS2 phenotype^9^ (Figure S2F-H). In fact, three MSI-H tumors in this atlas (SG00001, SG00002, PAT73458) exhibited high CNV scores, and were thus classified as CIN+. These exceptions are analogous to the hypermutated (HM) TA/TVA/MSS tumors (Figure 1F), suggesting that alternative pathways to CIN and HM will emerge in some cases.

In ST data, CNV inference provides tumor clone regions that spatially align with tissue domains annotated by histology as well as gene signatures enriched in dysplastic epithelium (Figure 2D). Specifically, ST CNV profiles cluster into major tumor clones based on similarity, while normal mucosa and stromal regions are aggregated into a background cluster (“S”; Figure 2D-H). Additionally, CNV inference identifies a tumor edge domain (“E”) resulting from a mixture of epithelium and surrounding stroma in ST microwells that dampens the CNV signal. These regions are useful for characterizing epithelial activity occurring at tumor borders such as epithelial-mesenchymal transition (EMT) and tumor cell invasion, as well as interactions with the tumor microenvironment (TME) including antigen presentation, lymphocyte exhaustion, cytotoxicity, and neutrophil recruitment (Figure 2D,G). Major clone regions can be ordered and labeled according to their inferred progression stages within each tumor, determined by a combination of CNV profile and mutational burden (TMB), as well as their relationships to one another (PAT71397 clone regions “1”, “2”, “3”; Figure 2E-H). Finally, ordering of CNV clones on an individual tumor basis is validated by tissue characteristics that trend with malignancy and progression, exemplified in PAT71397 by an increase in TMB, CD4+ T cell and suppressive T reg infiltration, and gene signature scores including iCMS2, stem, and CytoTRACE (Figure 2G).

### Multiregional somatic mutational profiles provide phylogeographical topology

Phylogenetic reconstruction provides insight into tumor progression from a snapshot in time, namely the moment of tumor resection and fixation. Our analysis revealed spatially informed clonal heterogeneity within tumors. We first called somatic mutations using bulk germline WES from each patient as a baseline (Methods: Somatic mutational profiling with LCM-WES). Results from 22 patients with LCM-WES exhibited observable phylogeny across three or more regions of interest (ROIs), deciphered using public, shared, and private mutations detected in spatially distinct tumor regions (Figure 3A-C; Figure S3A-W). Common and unique genetic alterations between ROIs allowed for the reconstruction of evolutionary relationships between tumor regions, which can follow one of several proposed evolutionary models^49–53^. Our data were consistent with three major models of tumor evolution (Figure 3D).

**Figure 3.**
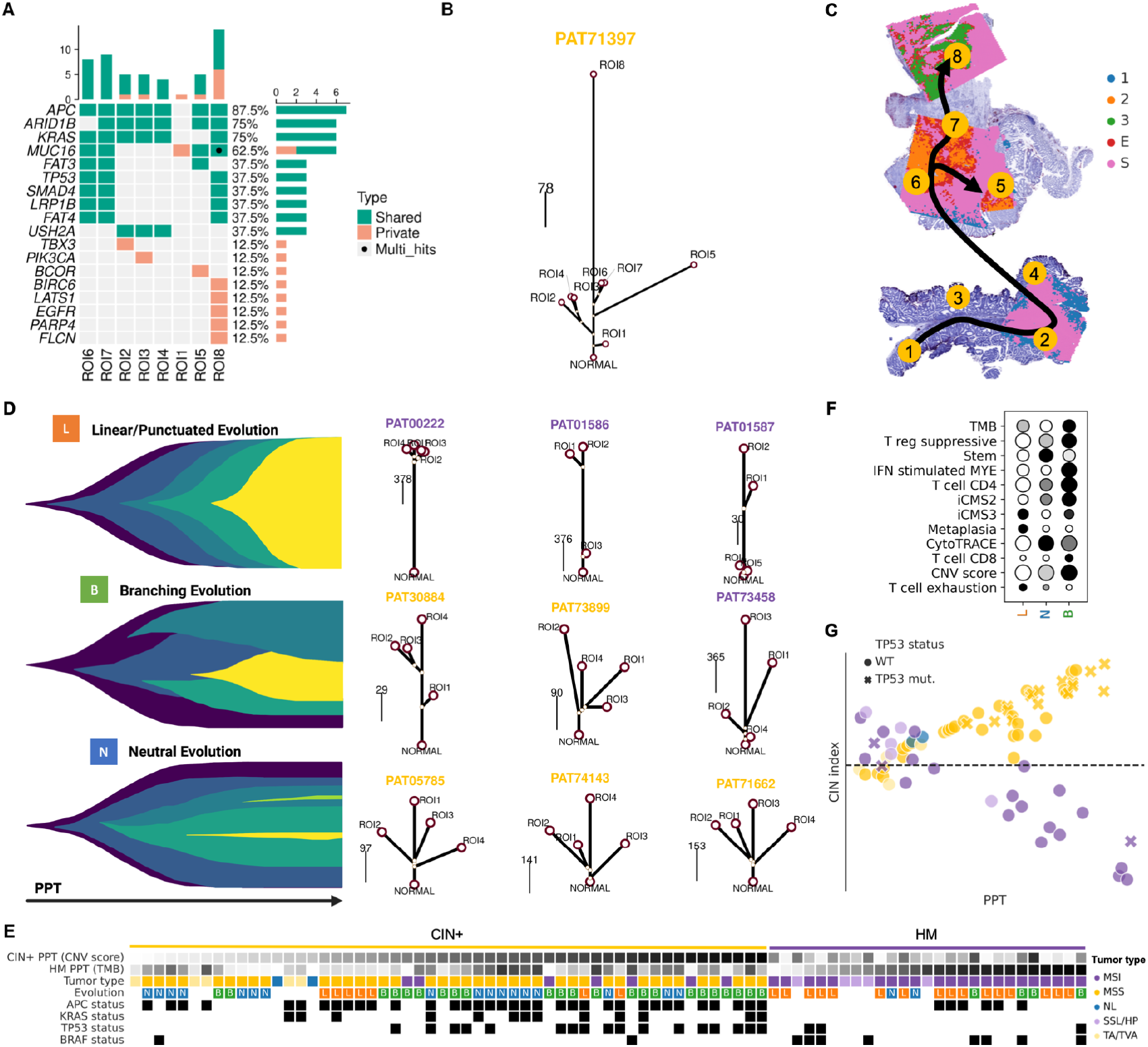
Multiregional somatic mutational profiles provide phylogeographical topology. (A) Oncoplot of detected driver mutations within spatially sampled LCM ROIs of PAT71397. (B) Phylogenetic tree for PAT71397. Length of branches are proportional to the number of shared or private somatic mutations in each LCM ROI. (C) Diagram of LCM ROIs in PAT71397 blocks, overlaid on CNV clone regions identified in ST. Black arrow represents inferred progression trajectory from CNVs and mutational phylogeny in B. (D) Diagram of observed modes of tumor evolution. Example phylogenetic trees from representative atlas samples shown to the right of each diagram. Patient ID colors represent tumor type (MMR status). (E) Tumor regions and their clinical and mutational metadata divided by class and ordered left-to-right by corresponding PPT (CNV score for CIN+, TMB for HM). (F) Summary of gene signatures across all tumor regions grouped by evolutionary mode from D. (G) CIN index versus PPT for tumor regions from E. Points are colored by tumor class, except pre-malignant and normal regions, which are colored according to tumor type as in Figure 1A-B. Points are colored by tumor class. Point shape corresponds to regions with detected *TP53* mutation.

Linear or punctuated evolution consists of stepwise increases in fitness that result in periodic “clonal sweeps” that replace the dominant makeup of the tumor^54^. These tumors typically exhibit low degrees of heterogeneity across regions with many shared or public mutations^51^. The neutral or “big bang” model of tumor evolution consists of many subclones with near-equal fitness due to early bursts of mutational events that persist throughout the life of the lesion^49^. Similarly, branching evolution is characterized by co-existing subclones that exhibit hierarchical relationships between one another.

When categorizing carcinomas in the atlas by phylogenetic structure, we observed that 8/22 tumors exhibited linear evolution characterized by many public mutations and low regional heterogeneity. Of the remaining 14 carcinomas, we classified six as neutral - effectively the opposite of linear, with few public mutations and high regional heterogeneity - and eight as branching. Interestingly, we classified the evolution of only 2/15 CIN+ tumors as linear. Conversely, linear evolution dominated the HM cohort (6/8 cases; Figure 3E; Figure S3X; Methods: Global PPT ordering and classification of tumor regions). Indeed, CIN+ signatures such as iCMS2, stem, CD4 T cells, and high CNV score are enriched in tumors with neutral and branching evolution, while HM markers including iCMS3, metaplasia, and T cell exhaustion are highly expressed by tumors undergoing linear evolution (Figure 3F). Despite a limited sample size, we speculate that hypermutation confers stronger and more protracted stepwise gains of clonal fitness, resulting in linear or punctuated evolution, whereas CIN promotes high-frequency, continual genetic alterations that yield minor, hierarchical differences between co-existing subclones in a branching or neutral evolutionary pattern^55–58^.

Given two distinct indicators of regional tumor progression, CNVs and somatic mutations, we next distinguished the major classes of colorectal tumors whose regional progression pseudotime (PPT) are best quantified by these indicators, respectively. Chromosomally unstable (CIN+) tumors, most likely MSS arising from the conventional adenoma pathway, should exhibit CNV profiles that reliably describe regional clonal relationships that recapitulate tumor progression topology. Hypermutated (HM) tumors, most likely arising from the serrated pathway, would conversely be MSI-H and chromosomally stable. Therefore, clonal progression is best described by regional TMB in the HM case.

To demonstrate these principles, we established PPT ordering and calculated a CIN index (quantifying the comparative degree of CIN and hypermutation) for tumor regions defined as a combination of LCM ROIs and CNV clones (Figure 3E,G; Methods: Global PPT ordering and classification of tumor regions). PPT ranking allowed for atlas-level integration and modeling of spatial information along a global indicator of progression from normal mucosa and pre-malignant adenoma to invasive adenocarcinoma. Advanced tumor regions (PPT > 0.4) with positive and negative CIN indices were classified as CIN+ and HM, respectively, uncovering spatial heterogeneity that describes transitions from MSI-H to CIN+ and MSS to HM. Importantly, we note that *TP53* mutations were more enriched than *APC* mutations in tumor regions with high CIN index, corroborating our observation that CIN emerges later in CRC development^1^ (Figure 3G; Figure S3Y).

### Cell-state deconvolution reveals pseudotemporal tissue dynamics

Using a previously published scRNA-seq dataset from 128 specimens consisting of CRCs, pre-malignant lesions, and adjacent normal mucosa^9^, we used non-negative matrix factorization^59^ (NMF; Figure S4A-B) to construct a consensus reference of 30 distinct cell states found in epithelial and stromal compartments and characterized by top-loaded genes and their associated pathway enrichment terms (Figure S4C-D). These cell states include normal mucosal cells such as tuft (TUF), enteroendocrine (EE1-2), and goblet (GOB), tumor-specific states derived from serrated lesions (SSC) and carcinomas (CRC1-4), and infiltrating immune populations including helper T cells (TL1), cytotoxic T cells (TL2), and neutrophils (MYE4), amongst others (Figure S4C-D; Table S3).

To calculate cell-state contributions to each ST pixel, we used reference (ref) NMF to extract consensus states from ST expression matrices, effectively inferring fractional abundance of these states in each ST microwell (Methods: refNMF cellstate discovery and deconvolution). Deconvolved cell-state fractions correlated with selected MxIF cell-type marker expression in registered serial tissue sections, as well as expression of literature-based cell-type and cell-activity gene signatures^60,61,9,62,10,32,11^ (Figure 4A-C; Table S2; Figure S4E-G; Methods: refNMF validation). For example, GOB, ABS (absorptive colonocytes), and CT (crypt-top colonocyte) enrichment aligns with high MUC2 IF staining in normal epithelium, MYE1 (M1 macrophages) coincides with CD11B staining, and CRC2/STM-rich regions (CIN+ tumor cells and stem-like cells) express PCNA and OLFM4 in PAT30884 (Figure 4A). The SSL, HTA11_08622_A, further validates refNMF states such as ABS, FIB2, and serrated-specific cells (SSCs) by their spatial distribution in the mucosa and submucosa layers and correspondence to MUC5AC and AQP5 IF staining (Figure 4B-C).

**Figure 4.**
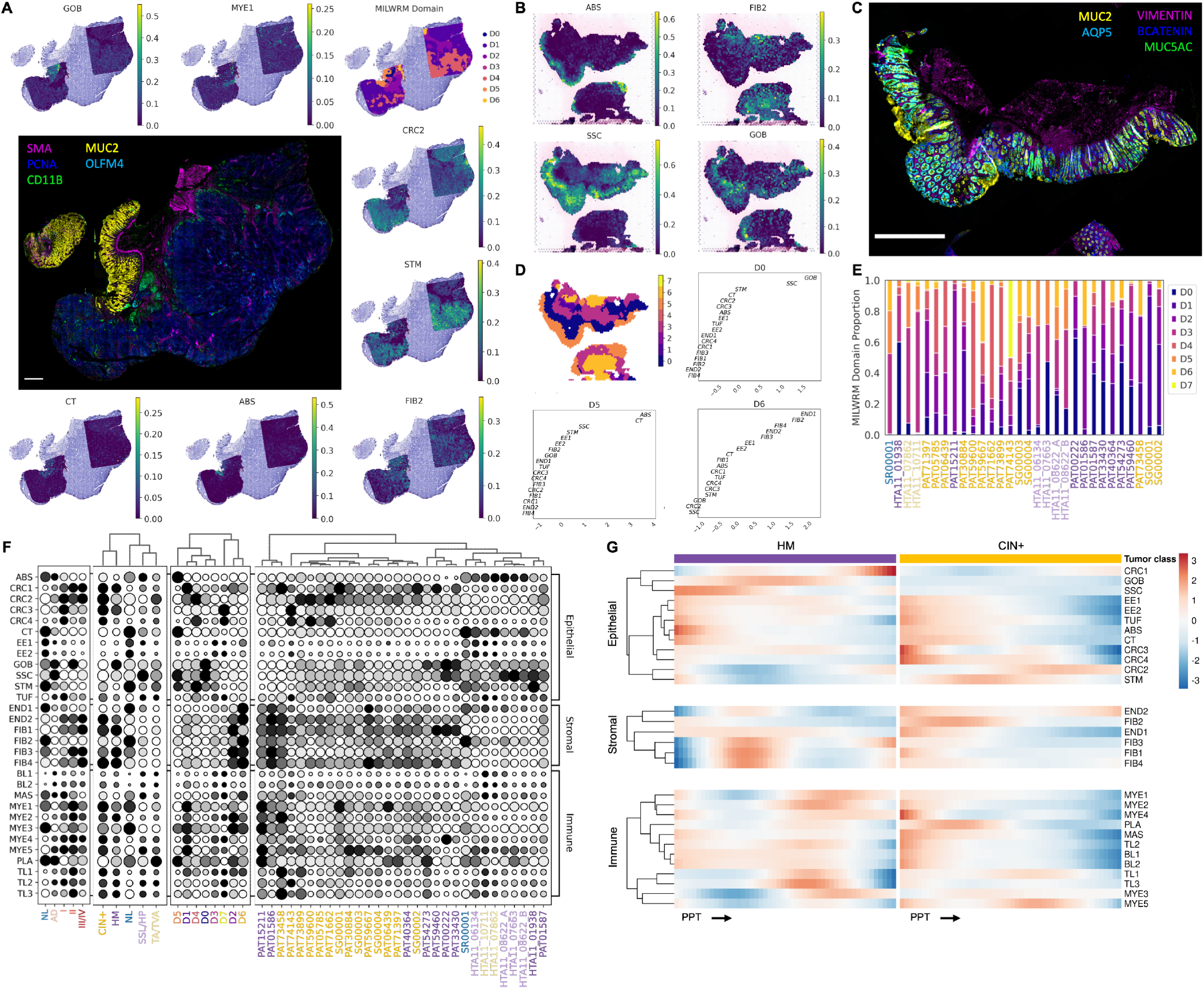
Cell-state deconvolution reveals pseudotemporal tissue dynamics. (A) Example whole slide MxIF surrounded by refNMF usages for seven cell states as well as MILWRM tissue domain projected onto PAT30884 histology. Scale bar 500 μm. (B) refNMF usages of normal absorptive colonocyte (ABS), normal fibroblast (FIB2), serrated-specific cell (SSC), and goblet cell (GOB) states for HTA11_08622_A. (C) MxIF image of HTA11_08622_A. Scale bar 500 μm. (D) MILWRM tissue domains for HTA11_08622_A, surrounded by top cell-state loadings for SSL (D0), normal epithelium (D5), and submucosa (D6) domains. (E) Proportions of MILWRM domains detected in ST from each patient. Patient ID colors represent tumor class. (F) refNMF states grouped by compartment and summarized across tumor stage, tumor class, MILWRM domain, and patient for all ST samples. Patient ID colors represent tumor class. (G) Heatmap of GAM fits for refNMF states in all ST tumor regions ordered by PPT for HM (left) and CIN+ (right) tumors. Color represents scaled expression within each tumor class.

Application of cell-cell interaction community reconstruction algorithms to ST data provided inaccurate results due to low spatial resolution and large distances between microwells. Focusing on pixel-based community detection, we employed refNMF usages as predictors for a MILWRM model to divide tissue into consensus domains based on cell-state makeup^63^ (Figure 4D). The MILWRM model yielded eight domains (D0-D8) that correspond to CIN+ epithelium (D4 - high in CRC2 and STM), normal mucosa (D5 - enriched in ABS and CT), and sessile-serrated epithelium (D0 - high in SSC and GOB), amongst others, all of which spatially align with regional histology (Figure 4A-E; Figure S4H).

When summarizing refNMF abundances across MILWRM domains and tumor classes, we can identify epithelial cell states enriched by unique populations (CRC1 = HM; CRC2 = CIN+) and gain a coarse understanding of microenvironmental makeup of each tumor (TL2 [cytotoxic] = SSL/HP/HM; MYE4 [neutrophils] and MYE5 [DCs] = CIN+; Figure 4F). Finally, we note that MILWRM domain D1 is characteristic of HM, while D4 is characteristic of CIN+ tumors (Figure 4E). Applying these approaches to ST data enables validated mapping of consensus tumor and non-tumor cell populations across the atlas, and specifically provided the spatial distributions of low-abundance cell states such as infiltrating immune cells.

Using PPT ordering from CNV scores and TMB in CIN+ and HM tumors, respectively, we tracked changes in refNMF states across a global indicator of tumor progression and built generalized additive models (GAMs) that ascribe statistical significance to cell-state dynamics during CRC evolution (Figure 4G; Table S4; Methods: Modeling expression dynamics along PPT). We observed a replacement of normal cells (ABS, CT, STM, TUF, and EE) with carcinoma-specific epithelial states (CRC1-4) following the transition from pre-cancer to CRC in both the CIN+ and HM pathways. Along CIN+ PPT, STM-dominated early lesions give rise to CRC2, while HM tumors progress from metaplastic SSCs to mucinous CRC (GOB and CRC1), likely due to the respective cells-of-origin of CIN+ and HM tumors^9^. Consistent with prior knowledge, we observed an increase in most infiltrating immune populations with development of HM tumors, but a striking decrease in immune states in the tumor epithelium of CIN+ CRCs, coincident with the strong increase in CRC2 abundance (Figure 4G). This observation, in the context of global PPT, suggests that an epithelial-intrinsic program emerges in CIN+ CRCs to exclude or evade the host immune system.

### Gene expression features of CIN+ CRCs predict immune exclusion

Immune exclusion correlates with poor patient outcomes in multiple cancer types and negatively predicts immune checkpoint inhibitor (ICI) response^24,64,65,19,66,28,62^.

Immune exclusion mechanisms provide potential therapeutic targets that may open the door for MSS CRCs to respond to ICI^67,68^. Biomarkers of immune exclusion may likewise act as predictors of the roughly 30 - 60 % of ICI non-responders in the MSI-H group that are currently immunotherapy candidates ^69^.

To further investigate the link between CIN+ tumor progression and immune exclusion, we extended GAM analysis to untargeted gene expression from ST atlas data. We identified genes with statistically significant dynamics within and between PPT trajectories for HM and CIN+ CRCs and grouped them by biological pathway or function (Figure 5A; Table S4). Specifically, we identified genes that were significantly elevated in late CIN+ PPT, which formed a functional module for extracellular matrix (ECM) signaling and organization (*DDR1, TGFBI*, and *PAK4*). These genes originate from epithelial cells, which implicated a mechanism of microenvironmental modulation by the tumor itself. Each of these genes has also been shown to associate with immune exclusion in several solid tumor types, providing an interesting, concerted signature that potentially creates an immune-tolerant environment in CIN+ CRC^70,26,27,71^.

**Figure 5.**
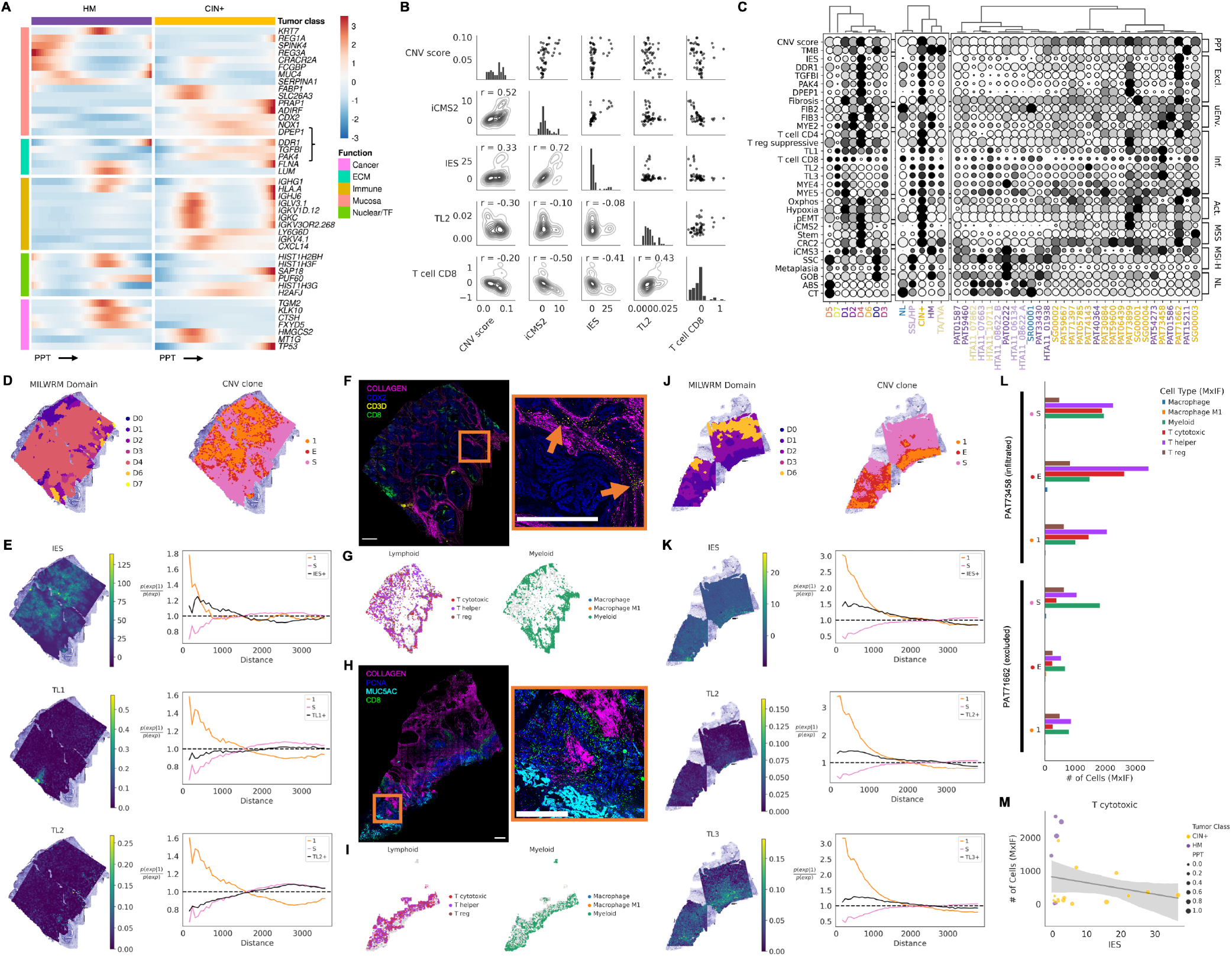
Gene expression features of CIN+ CRCs predict immune exclusion. (A) Heatmap of GAM fits for top genes summarized across all ST tumor regions. Color represents scaled expression within each tumor class. Genes are grouped by biological function. Bracket denotes IES genes. (B) Pairwise Pearson correlations between progression indicators (CNV score and iCMS2), IES, cytotoxic T cell refNMF state (TL2), and CD8 T cell gene signature in all CIN+ tumor regions. (C) Genes, gene signatures, and refNMF states grouped into pseudotime indicators (“PPT”), immune exclusion markers (“Excl.”), microenvironmental cells (“uEnv.”), infiltrating immune cells (“Inf.”), tumor activity (“Act.”), and epithelial-specific markers of MSS, MSI-H, and normal mucosa summarized by MILWRM domain, tumor class, and patient for all ST samples. Patient ID colors represent tumor class. (D) PAT71662 ST with annotated tissue domains from MILWRM (left) and CNV clone regions (right). (E) Expression overlay and spatial co-occurrence analysis for IES, helper T cells (TL1), and cytotoxic T cells (TL2) in PAT71662. Line plots at right indicate the conditional probability of high signature or cell-state expression as a function of distance from CNV clone 1 microwells (Methods: Spatial co-occurrence analysis from ST). (F) PAT71662 MxIF showing collagen, CDX2 (marking MSS epithelium), and lymphocytes (CD3 and CD8). Inset highlights CD3/CD8+ cells sequestered to stroma. Scale bars 500 μm. (G) Centroids of segmented single cells from PAT71662 MxIF plotted in whole-slide space, split into lymphoid and myeloid compartments. (H) Same as F for PAT73458. PCNA and MUC5AC mark tumor epithelium. Inset highlights CD8+ cells invading epithelium. Scale bars 500 μm. (I) Same as in G, for PAT73458. (J-K) Same as in D-E, for PAT73458. TL3 represents *γδ*IELs. (L) Census of infiltrating immune cells in PAT71662 (bottom) and PAT73458 (top) from G and I summarized by CNV clone region (Methods: MxIF immune-exclusion analysis). (M) Number of infiltrating CD8+ T cells detected in MxIF plotted against IES score for all tumor regions. Points are colored by tumor class and sized according to PPT ranking.

Additionally, we observed a significant increase in *DPEP1* expression with CIN+ PPT, a gene which was also highly enriched in the CRC2 epithelial state (Figure 5A; Figure 4H; Figure S4C-D). We added *DPEP1* to the signature with *DDR1, TGFBI,* and *PAK4* for analysis of immune exclusion in this atlas as it is implicated in neutrophil recruitment, which could lead to lymphocyte exclusion in the TME ^72,73^, and has been shown to be secreted by tumor cells in extracellular vesicles, thus offering a promising circulating biomarker in CRC patients ^74^.

These four genes proved to be highly co-expressed by tumor epithelium (Figure S5A-B), and when combined into an Immune Exclusion Signature (IES), exhibited spatial correlation with iCMS2 and CNV score, and were enriched in the CIN+-specific MILWRM domain D4, corroborating prior studies^23,25,10^ (Figure 5B-C). Surveying other gene signatures, we identified additional programs that coincided with IES such as pEMT and oxidative metabolism, which are hallmarks of advanced cancer and upregulated during tumor progression, validating the emergence of IES in late PPT (Figure 5C; Figure S5B). Moreover, fibrosis and hypoxia signatures correlated with IES at the patient-level, lending credence to the involvement of constituent genes in ECM signaling and regulation as these characteristics are implicated in microenvironmental immunosuppression^70,26,27,71,75^ (Figure 5C; Figure S5D). Most importantly, this four-gene IES had a negative correlation with cytotoxic T cells (TL2) and with the CD8 T cell signature score across all tumor regions in the dataset, indicating its value as a predictor of immune exclusion^62^ (Figure 5B).

An example from this atlas of immune-excluded CRC is PAT71662, which contains a single major CNV clone region that delineates the epithelial compartment in ST (clone region “1”; Figure 5D). We note that the majority of the area of this MSS tumor belongs to MILWRM D4, and is correspondingly high in CRC2, stem, fibrosis, and oxidative metabolism (Figure 5C-D). Spatial co-occurrence analysis revealed that this tumor exhibited enrichment of the epithelial-intrinsic IES in the tumor core, while excluding T lymphocytes from the CNV clone 1 region (Figure 5E). We confirmed the exclusion of CD8+ cells using spatially registered MxIF data, demonstrating how immune infiltrates are sequestered to the collagen-rich stroma (Figure 5F-G). Conversely, immune-infiltrated tumors such as PAT73458 exhibited much lower epithelial expression of IES while clear infiltration of cytotoxic (TL2) and *γδ* (TL3) T cells is seen in ST and MxIF data (Figure 5C,H-K).

We next expanded immune-exclusion analysis using MxIF data to validate ST findings. Following single-cell segmentation^76^, we identified immune cell subsets by the presence of various marker protein stains quantified in each cell. We then masked the MxIF slides with CNV clone regions so we could enumerate the abundance and distribution of infiltrating immune cells in each major tumor compartment (Methods: MxIF immune-exclusion analysis). We note that immune-excluded PAT71662 has more immune cells in its stroma (“S”) than tumor epithelium (“E” and “1”), while the infiltrated tumor, PAT73458, exhibits not only higher levels of cytotoxic (CD3+/CD8+) and helper (CD3+/CD4+) T cells overall, but also a larger proportion of all infiltrating immune cells in the tumor edge and tumor core regions (Figure 5F-I,L). Extending these analyses to the entire spatial atlas, we observe a negative correlation between IES and T cells (CD8+ and CD4+) in tumor clone regions as identified by high-resolution MxIF imaging (Figure 5M; Figure S5C).

In HM/iCMS3 tumors with low IES, we observed a corresponding increase in other microenvironmental cell states that have been implicated in immune tolerant microenvironments, suggesting that HM CRCs follow an alternative path to immune evasion and potential ICI non-response compared with CIN+ CRCs. Cancer-associated fibroblasts (CAFs) expressing *FAP* and *CXCL12* (FIB3), and *SPP1+* myeloid cells (MYE2) have been shown to foster an immunosuppressive niche in CRC and other solid tumors such as pancreatic cancer^22,77,78^ (Figure 5C; Figure S5E). Moreover, *CXCL14+* CAFs (FIB2) are suspected to counteract the immune-silencing effect of *CXCL12+/FAP+* fibroblasts and are more prominent in MSI-H/iCMS3/HM tumors^28^. Since these three microenvironmental cell states emerge in immune infiltrated tumors (Figure 5C) and seem to trend positively with regional HM progression (Figure 6A), we hypothesize that this emergent struggle between immune evasive and immunogenic TMEs represents intracellular signaling complexity in late-stage CRC, and that tipping the scales of such interactions may explain the subset of MMR-deficient HM tumors that do not respond to ICI therapy^78^.

**Figure 6.**
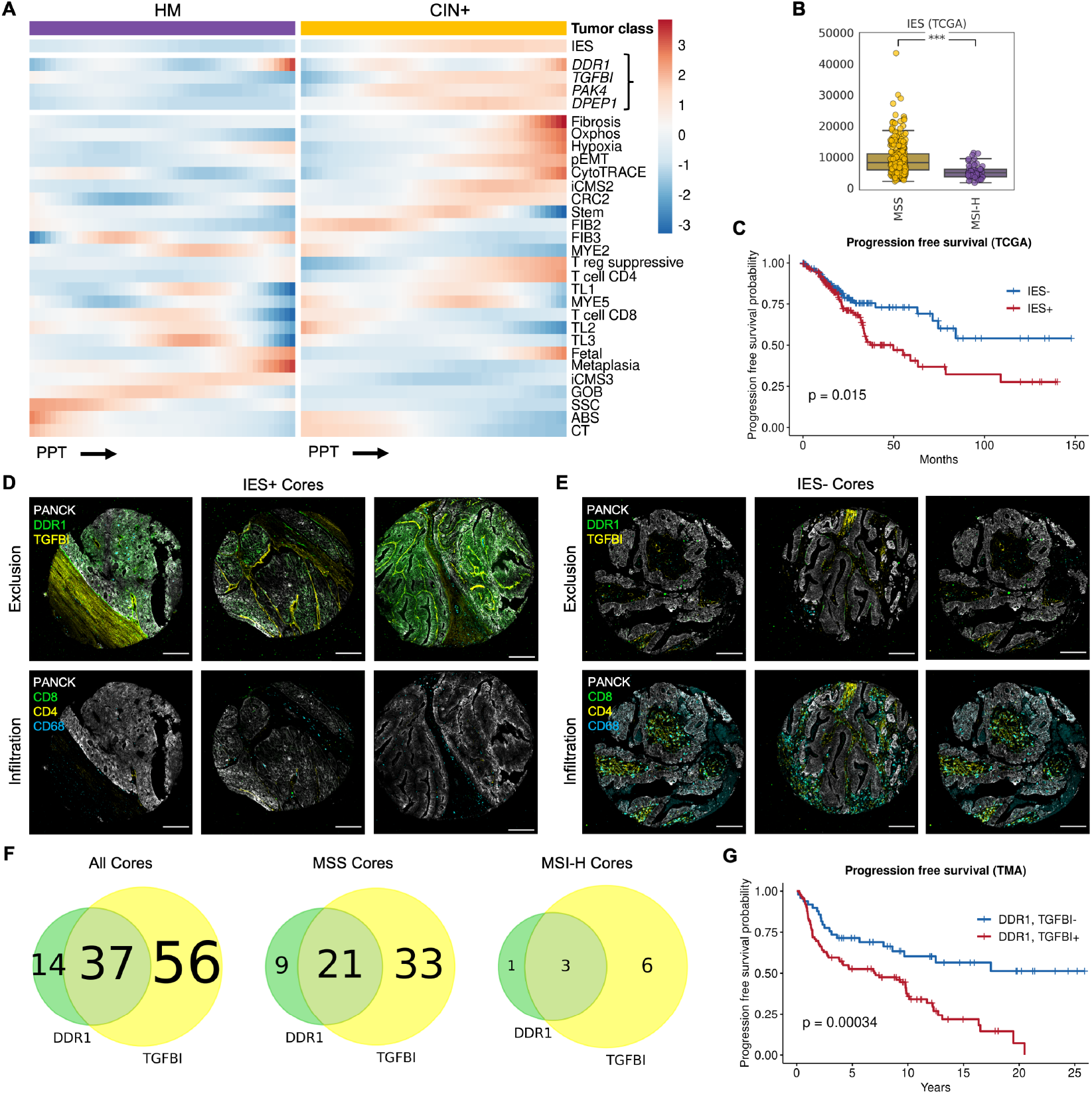
IES trends with tumor progression and predicts poor patient outcomes. (A) Heatmap of GAM fits for genes, gene signatures, and refNMF cell states summarized across all ST tumor regions. Color represents scaled expression within each tumor class. Bracket denotes constituent genes in IES. (B) Boxplots of IES scores in TCGA COAD and READ samples, stratified by MMR status (MSS *n* = 301; MSI-H *n* = 45). Student’s T-test with Bonferroni correction yielded p = 6.04 × 10^−8^ (C) Kaplan-Meier PFS curves for TCGA COAD and READ samples from B with high (+) and low (-) IES scores. (D) MxIF images showing epithelial (top) and immune (bottom) markers from representative IES+ cores from the CRC TMA. Scale bars 100 μm. (E) Same as in D, for representative IES-cores. (F) Venn diagram of the number of CRC TMA cores with high-scoring DDR1 and TGFBI IHC staining (left; total *n* = 163), and stratified by MMR status (middle, *n* = 86; right, *n* = 22). (G) Kaplan-Meier PFS curves for CRC TMA cores with high (+) and low (-) IHC staining of both DDR1 and TGFBI.

### IES trends with tumor progression and predicts poor patient outcomes

To further explore the link between tumor progression and immune exclusion, we modeled scoring of CIN+ epithelial-intrinsic IES, HM-enriched FIB2 and FIB3 CAFs and MYE2 macrophages, as well as infiltrating immune cell states and cancer progression signatures against PPT for all tumor regions across the atlas, confirming that IES is enriched in CIN+ tumors and trends proportionally to CNV-informed PPT, iCMS2 and CRC2 enrichment (Figure 6A; Table S4; Methods: Modeling expression dynamics along PPT).

We next set out to validate this expression signature as a predictor of immune exclusion by highlighting its translational utility in predicting patient outcomes in external cohorts with larger sample sizes. The IES score was significantly enriched in MSS (CIN+) vs. MSI-H (HM) TCGA COAD and READ samples (p = 6.04 × 10^−8^; Figure 6B; Figure S6A), consistent with the distinct immunosuppressive mechanisms between the two tumor subtypes. High IES expression yielded a statistically significant drop in progression-free survival (PFS) for patients with high-scoring tumors compared to those with low-scoring tumors from the entire TCGA cohort (p = 0.015; Figure 6C) as well as the subset of MSS tumors in TCGA (p = 0.035; Figure S6B). Furthermore, only *TGFBI* exhibited a similar reduction in PFS by itself (Figure S6C-D), indicating that the epithelial-intrinsic IES score has prognostic value in identifying immune-cold CRCs with poor disease-free survival, and that the aggregate signature is more informative than the sum of its parts.

Finally, to increase the translational value of this signature, we investigated whether protein expression corroborated results obtained by gene expression in CRC such that immunohistochemistry (IHC) can be reliably used to predict immune exclusion. From IHC staining of a tumor microarray (TMA) consisting of 163 colorectal adenocarcinoma samples, we observed higher overall DDR1 and TGFBI protein expression in MSS versus MSI-H tumors, consistent with findings above (Figure 6D-F). Survival analysis on this TMA IHC cohort revealed significantly lower PFS (p = 0.00034; Figure 6G) and overall survival (OS; p = 0.0011; Figure S6E) for tumors with high IHC staining for both DDR1 and TGFBI, corroborating our TCGA query. Once more, this combination of IES markers yielded a statistically greater stratification of patient survival than the individual proteins (Figure S6F-G), confirming the utility of this expression signature as a prognostic indicator in both mRNA and protein assays.

## Discussion

Evolution of spatially resolved tumor clones in human CRC presents an opportunity to use spatial technologies to investigate cellular and microenvironmental heterogeneity along a trajectory of cancer progression. Whereas the probability of any particular polyp progressing to malignancy is inherently low^5,79^, this spatial strategy of mapping evolution in pre-selected carcinomas with less advanced components offers a glimpse into the biology of malignant transitions that can inform diagnostic stratification and early intervention^49,30,16,80,81^. The phylogeographical atlas presented herein comprises spatially resolved genomic, transcriptomic, and protein profiling of 31 patients spanning normal mucosa, pre-cancerous lesions, and invasive adenocarcinoma. Spatial assays in each of these molecular domains recapitulated findings related to TA/TVA and SSL/HP progression to MSS and MSI-H CRCs from initial *WNT* activation and gastric metaplasia, respectively, to more advanced iCMS states^9,10^.

Using CNV and LCM-WES analysis, we classified tumors as CIN+ or HM, and confirmed that the latter group displays a cytotoxic immune microenvironment while the former are more likely to be immune-cold^7,4,25^. While previous work documented a mechanism by which *APC* loss-of-function leads to aneuploidy and chromosomal instability ^41,43,42^, our multi-modal CNV analysis supports that most CIN occurs as a late-onset characteristic within the adenomacarcinoma sequence, likely following *TP53* loss and transition to malignancy^82–85^. In this regard, some MSI-H tumors (SG00001, SG00002, PAT73458) gain CIN through APC-independent mechanisms. Likewise, some MSS cases display an HM phenotype (HTA11_01938, PAT15211), likely driven by mutations in proofreading polymerases such as *POLE* and *POLD1* ^37^. Phenotypic manifestations are more consistent with CIN+ and HM classifications than microsatellite status, exemplified by MSI-H/CIN+ tumors transitioning to a stem-like, iCMS2 state^9^.

Multiregional somatic mutational profiling matched CNV phylogeography in CIN+ CRCs, provided high-confidence PPT for HM tumors, and allowed us to stratify patients based on evolutionary dynamics. In this atlas, we observed CRCs that underwent neutral or hierarchical evolution, exhibiting high degrees of regional heterogeneity, as well as tumors that displayed linear or punctuated evolution with many public driver mutations and relatively low clonal divergence ^49,51^.

We found that neutral and branching evolution dominated CIN+ tumors, while the majority of HM CRCs exhibited linear or punctuated evolution. Although our dataset lacks sufficient sample size to draw confident conclusions, we speculate that this dichotomy of evolutionary dynamics stems from the discrepancy between somatic mutation rate in HM CRCs and genomic structural alterations due to CIN, resulting in large, stepwise increases in clonal fitness and smaller, more frequent subclonal branching, respectively^55–58^. Furthermore, tumors driven by distinct mutational processes may recruit unique TMEs that further restrict clonal evolution, such as distinct immune cells or CAFs that suppress or promote tumor progression^86,77,87,24^. Nevertheless, observed evolutionary dynamics in this atlas serve to characterize overall tumor heterogeneity and enable pseudotemporal placement along a global progression trajectory.

Generalized additive modeling (GAM) along global PPT elucidated tumor and microenvironmental programs such as altered metabolism that drive or result from tumor progression^88,89^. Of specific interest are gene programs that modulate tumor-microenvironment interactions. We focused on an immune-exclusion mechanism specific to CIN+ tumors with an iCMS2/stem-like epithelial phenotype marked by the CRC2 refNMF state. Previous work from our group and others has demonstrated a transition into a stem-like tumor state as a function of progression, resulting in immunosuppression^90,9^. PPT models enabled the unbiased identification of gene programs relevant to tissue dynamics along tumor development, and allowed us to assemble an epithelial-intrinsic Immune Exclusion Signature (IES) defined by expression of *DDR1, TGFBI, PAK4,* and *DPEP1.* This aggregated signature is highly informative, as these genes have been implicated separately as immune modulators but not always in the context of CRC^26,91,92,71^.

*DDR1* has been reported to align collagen fibers in a way that excludes T cells from the breast TME ^27^. These findings, coupled with *DDR1*’s role in fibrotic kidney disease and breast cancer, validate the observation that IES coincides with fibrosis and hypoxia in advanced CIN+ CRCs and suggest that microenvironmental signaling and collagen remodeling in the ECM are involved directly in immune evasion^93,94^. *TGFBI* is associated with an immunosuppressive microenvironment in ovarian cancer where it is released from macrophages, but mechanistic details are lacking^95^. *DPEP1* is an endothelial adhesion receptor for neutrophils and monocytes during inflammation, and while a direct role for *DPEP1* in CRC immune evasion has not been reported, it has been shown that neutrophil infiltration can lead to T cell exclusion in the colon^72,73^. We note that neutrophil infiltration increased with PPT in our dataset, highlighting the importance of *DPEP1* in CRC progression.

A confluence of these immunomodulatory processes has also been observed in a recent tumor microbiome study, where advanced, microbially-infiltrated CRC regions are aneuploid, hypoxic, and immunosuppressive in a neutrophil-dependent manner^96^. Altogether, these molecules play distinct yet complementary roles in immune exclusion that will be the focus of future studies.

HM tumors, on the other hand, exhibited a stromal dichotomy of immune surveillance escape driven by *CXCL12+* CAFs and *SPP1+* myeloid cells^78^, and *CXCL14+* iCAFs that have been shown to be MSI-H-specific and immunogenic^28^. Importantly, we note that the prevalence of these tumor-immune interaction networks trends directly with HM PPT, suggesting a link to CRC development analogous to IES in the CIN+ cohort.

The immune exclusion biomarkers identified herein carry several clinically relevant implications for detection, prognosis, and potential treatment of CRC. IES genes and their protein products in aggregate offer a robust prognostic indicator of progression-free survival in CRC. Additionally, this expression signature may stratify patients by ICI response potential: identifying ICI responders in the CIN+/MSS cohort as well as ICI non-responders in the HM/MSI-H group. Moreover, these biomarkers may act as potential targets for adjuvant therapy to ICI, depending on tumor stage and CIN status, as the breakdown of immune evasive mechanisms could expose tumors to more effective immunotherapy.

Here we present a rich, spatially resolved dataset comprised of genetic, transcriptomic, and proteomic layers of molecular information. Integrated analyses herein aimed to map recurring cell states and signatures across patient-specific phylogeographical landscapes to uncover global pseudotemporal dynamics of tumor progression from a snapshot in time. However, there is yet much more to be learned regarding CRC progression, and these data will therefore prove to be a valuable resource in the further characterization of tumor-microenvironmental co-evolution.

## Supporting information

Key Resources Table

## ACKNOWLEDGEMENTS

The authors wish to thank the study participants and other contributing investigators including Vito Quaranta, Jake Hughey, and Sarah Groves. We apologize in advance to those we have failed to acknowledge due to space constraints. This study was supported by the Human Tumor Atlas Network grant U2CCA233291 (to R.J.C., K.S.L., M.J.S., and C.L.S.), R35CA197570 and P50CA236733 (to R.J.C.), R01DK103831 (to K.S.L.), K07CA122451 (to M.J.S.), F31DK127687 (to P.N.V.), and P30CA068485 (to Vanderbilt-Ingram Cancer Center). Polyp RNA-seq funding was provided by Janssen (to M.J.S.). Cores used by this study included TPSR (U24DK059637), DHSR, the CHTN (UM1CA183727), and VANTAGE. A portion of the participants were studied as the result of resources and the use of facilities at the Veterans Affairs Tennessee Valley Healthcare System. R.J.C. acknowledges the generous support of the Nicholas Tierney GI Cancer Memorial Fund.

## Author contributions

Conceptualization, C.N.H., M.J.S., R.J.C., and K.S.L.; data curation, C.N.H., A.J.S., F.R., J.W., H.K., B.C., M.A.R.-S., E.T.M., P.N.V., J.T.R., M.K.W., M.J.S., and K.S.L.; formal analysis, C.N.H., E.T.M., M.A.R.-S., D.A., Y.W., S.E.G., B. C., N.S., J.L.D., A.R., Q.L., and K.S.L.; investigation, C. N.H., A.J.S., M.K.W., C.L.S., M.J.S., R.J.C., and K.S.L.; methodology, C.N.H., A.J.S., E.T.M., Y.W., D.A., S.V., H.K., and K.S.L.; project administration, C.N.H., A.J.S., E.T.M., J.T.R., Q.L., M.K.W., M.J.S., R.J.C., and K.S.L.; resources, M.K.W., J.T.R., Q.L., M.J.S., R.J.C., and K.S.L.; software, C.N.H., H.K., B.C., J.S., E.T.M., Q.L., and K.S.L.; supervision, M.K.W., R.J.C., M.J.S., and K.S.L.; validation, H.K. and C.N.H.; visualization, C.N.H., Y.W., D.A., B.C., R.J.C., M.J.S., and K.S.L.; writing – original draft, C.N.H., R.J.C., and K.S.L.; writing – reviewing and editing, C.N.H, A.J.S., F.R., E.T.M., M.A.R.-S., J.W., J.S., Y.W., D.A., S.E.G., B.C., H.K., P.N.V., J.L.D., A.R., N.S., S.V., M.K.W., J.T.R., C.L.S., Q.L., M.J.S., R.J.C., and K.S.L.

## Data and code availability

Data have been deposited to the HTAN Data Coordinating Center Data Portal at the National Cancer Institute: https://data.humantumoratlas.org/ (under the HTAN Vanderbilt Atlas).

## Methods

### Sample procurement

These specimens were procured through the collaborative human tissue network (CHTN) as formalin-fixed, paraffin-embedded (FFPE) tissue blocks with accompanying pathology reports. Patient metadata is available in Table S1.

### Visium ST sample handling

Regions of interest (ROIs) for ST were chosen based on histological annotation of FFPE blocks, targeting tumor areas with morphology indicative of various stages of malignancy, and transition points between them. Tissue sections were cut and trimmed (if necessary) into 6.5mm x 6.5mm capture areas of 10X Genomics Visium FFPE spatial gene expression slides (Key Resources Table). Serial tissue sections were collected simultaneously for whole-slide MxIF staining (Methods: Multiplex immunofluorescence (MxIF) imaging) and laser capture microdissection (Methods: Somatic mutational profiling with LCM-WES).

Visium FFPE spatial gene expression slides were temporarily coverslipped, stained with hematoxylin and eosin (H&E; Key Resources Table), and brightfield imaged at 20X objective prior to tissue permeabilization, probing, and library prep according to the 10X Genomics protocol. Sample libraries were sequenced on an Illumina NovaSeq targeting 125M reads per capture area. Resulting sequencing data were aligned using 10X Genomics Space Ranger software version 1.3.0 (Key Resources Table).

### Visium TMA building

A subset of FFPE blocks was chosen for building tissue microarrays (TMAs), where three to five 1mm punches were collected and arrayed in a 3 x 3 format for sectioning into 6.5mm x 6.5mm capture areas of 10X Genomics Visium FFPE spatial gene expression slides (Key Resources Table). Adjacent 1mm punches to each TMA region of interest (ROI) were collected in tubes for direct DNA extraction and whole-exome sequencing (WES) library preparation, providing analogous molecular information to LCM-WES data (Methods: Somatic mutational profiling with LCM-WES).

### Multiplex immunofluorescence (MxIF) imaging

A cyclic staining, imaging, fluorophore inactivation protocol for multiplexed protein imaging was employed as shown previously^33^. A panel of 33 antibodies was used for staining, as detailed in Key Resources Table.

Virtual H&E stains were generated from autofluorescence (AF) images for each block, and are used to orient Visium data to whole-slide tissue morphology following spatial registration (Methods: Spatial registration).

### Spatial registration

We developed a custom Python plugin for napari, a multidimensional viewer for biological images^97^, which allowed us to perform affine transformation and scaling on Visium brightfield images in order to spatially align morphology features of the tissue with wholeslide MxIF. The `napari` plugin exports affine matrices and final image sizes, which can be applied directly to MxIF to align pixels with Visium ST microwells, or applied in reverse to Visium spots to cast them into whole-slide space on top of MxIF images.

Registration of LCM-WES with ST was performed manually by creating masks for each LCM ROI in the 10X Genomics Loupe Browser that were used to subset Visium microwells to LCM ROIs for downstream analysis.

### Gene signature scoring

Gene expression signatures were curated from the literature to interrogate cellular identity and activity in scRNA-seq and ST samples^60,61,9,62,10,32,11^. Lists of genes from Table S2 were passed to the `scanpy` function `score_genes`, with default parameters. There were two exceptions:

- “CytoTRACE” scores were calculated using the Cyto-TRACE R package version 0.3.3^61^
- When scoring “iCMS2” and “iCMS3” tumor epithelial signatures from Joanito, *et al.,* genes from the “Up” list were scored against genes from the “Down” list from each respective iCMS classification by restricting the `gene_pool` parameter of the `score_genes` function to the combined “Up” and “Down” gene lists^10^

### CNV inference from ST and scRNA-seq

We used the `infercnvpy` Python package to infer somatic CNVs from scRNA-seq and ST gene expression^98^. Analysis was performed separately on scRNA-seq and ST. For scRNA-seq analysis, we used all normal epithelial and stromal cells from each patient, as labeled in^9^, to provide a normal background for calling CNVs in malignant cells derived from the same patient. In ST, we grouped Visium microwells by patient and used stromal regions and adjacent normal epithelium, manually annotated in the 10X Genomics Loupe Browser, to provide a normal background for calling CNVs in tumor regions. For patient samples lacking sufficient stromal/normal surface area (mostly pre-malignant polyps and TMAs), we grouped multiple patients with the normal human Swiss roll sample (SR00001), which provided a chromosomally-stable reference for inferCNV analysis. Grouped samples included SR00001 (NL), HTA11_01938 (TVA), WD33469 (TMA), WD33473 (TMA), and WD33474 (TMA).

### CNV calling from WES and WGS

FFPE curls were treated with the truXTRAC FFPE total NA kit from Covaris (Key Resources Table) to extract DNA prior to library preparation. Whole-genome (WGS) libraries were prepared using a modified Twist Bioscience Human Genome Panel (Key Resources Table) and sequenced on an Illumina NovaSeq, targeting 50X coverage genome-wide. Whole-exome libraries were prepared using the Twist Bioscience Human Comprehensive Exome Panel (Key Resources Table) and sequenced on an Illumina NovaSeq, targeting 50X coverage exome-wide.

Somatic CNVs were called following `GATK4` Best Practices workflow and annotated with `GATK4` Funcotator data sources v1.6. We used the `CNVkit` Python package to summarize and visualize somatic CNVs in WES and WGS samples^99^.

### Somatic mutational profiling with LCM-WES

We performed laser capture microdissection (LCM) on FFPE sections serial to Visium ST samples (Methods: Visium ST sample handling) using the Arcturus XT LCM system from Thermo Fisher Scientific. Circular ROIs 1.5mm - 2.0mm in diameter were collected from spatially distinct tissue regions, targeting abnormal tumor epithelium with morphology indicative of various stages of malignancy, and transition points between them. For samples lacking adjacent normal colon biopsies or whole-blood samples for bulk WES, we dissected additional ROI(s) from adjacent normal epithelium or stroma present in the FFPE sections to use as germline reference for mutation calling.

We extracted DNA from dissected FFPE ROIs using the Arcturus PicoPure DNA Extraction Kit from Applied Biosystems (Key Resources Table). Alternatively, FFPE cores from Visium TMAs were treated with the truXTRAC FFPE total NA kit from Covaris (Key Resources Table) to extract DNA prior to library preparation (Methods: Visium TMA building). Whole-exome libraries were prepared using the Twist Bioscience Human Comprehensive Exome Panel (Key Resources Table) and sequenced on an Illumina NovaSeq, targeting 50X coverage exome-wide.

FASTQ reads were trimmed to remove adapter sequences using `Cutadapt` v2.10. Quality control on both raw reads and adapter-trimmed reads was performed using `FastQC` v0.11.9. The reads were then aligned to the human reference genome `hg19` using BWA v0.7.17. Duplicated reads were removed and the alignments were refined with `GATK4` Mark Duplicates and Base Quality Score Recalibration tools. Somatic variants were called using `GATK4 Mutect2` in “normal-tumor” paired mode and annotated with `ANNOVAR` ^100^ (2019/Dec/05 version).

### Phylogenetic tree construction from LCM-WES

Mutation Annotation Format (MAF) output files from WES alignment and mutation calling (Methods: Somatic mutational profiling with LCM-WES) were processed with the `maftools` R package to generate oncoplots and summarize mutations within and between samples^101^. We then used the `MesKit` R package to generate phylogenetic trees for each patient^102^.

### Global PPT ordering and classification of tumor regions

Following spatial registration of ST and LCM-WES, we divided tumors into areas of overlap between ST CNV clones (non-”E” and non-”S” regions) and LCM ROI masks (referred to as “tumor regions”), in order to provide paired mutational and CNV information for each region (Methods: Spatial registration). Tumor regions containing less than 30 microwells were disposed for reliable summarization of cell states, genes, and gene signatures. Pre-malignant tumors (TA/TVA and SSL/HP) were simply subset to their major CNV clone region, as their WES data was collected in bulk due to small size of the lesions preventing LCM analysis. The normal colon specimen (SR00001) was also subset to its epithelial area using the two CNV clone regions detected. Additionally, we removed HTA11_01938 from tumor region ordering due to extreme hypermutation likely resulting from compromised DNA excision repair and proofreading machinery (Figure 1F). Exclusion of this outlier prevents skewing of cell states and gene expression by a pre-malignant polyp assigned to late HM PPT.

Resulting tumor regions comprising groups of ST microwells sub-divided each macro tumor into smaller, epithelial-only regions which could be summarized based on average expression of genes, gene signatures, and cell states. We then calculated CIN+ PPT, defined as CNV score scaled between 0 and 1 using `sklearn.preprocessing.MinMaxScaler`^103^ across all tumor regions. Likewise, HM PPT was calculated as TMB scaled between 0 and 1 using `sklearn.preprocessing.MinMaxScaler`^103^ across all tumor regions. Next, we created a quantitative “CIN index” for each tumor region, calculated as the difference between CIN+ PPT and HM PPT. In this way, tumor regions with a CIN index greater than zero are more chromosomally unstable than hypermutated, and regions with a negative CIN index are conversely more hypermutated.

We then classified these CNV clone-LCM ROI overlap regions as CIN+ or HM based on their tumor type (MMR status and pathological annotation) coupled with their CIN index. All tumor regions with PPT values less than 0.4 were assigned to tumor classes according to the tumor type they were derived from (NL, TA/TVA and MSS = CIN+; SSL/HP and MSI-H = HM). Seven late-stage (PPT > 0.4) MSI-H tumor regions had CIN indices > 0.0 (2/4 regions from SG0001, 2/3 regions from SG00002, and 3/4 regions from PAT73458), and were thus labeled CIN+ for downstream GAM modeling and analysis (Figure 3E,G; Figure S3X-Y).

We assigned CIN+ tumor class to SG00001, SG00002, and PAT73458 for the purposes of patient-level summarization and downstream analyses (Figure 4F-G; Figure 5C), as the majority of the tumor area of these MSI-H CRCs had transitioned to CIN+. Additionally, one tumor that was excluded from tumor region analysis above due to low number of ST microwells available for each CNV clone-LCM ROI overlap region, PAT15211, was categorized as HM due to high TMB (low CIN index). All other analyses were performed at the tumor region level and employed classifications described above (Figure 4H; Figure 5A-B; Figure S5A; Figure 6A). Finally, we ordered all tumor regions along PPT according to their tumor class (Figure 3E; Figure S3X). These rankings provided a pseudotemporal basis for GAM fitting along two tumor growth trajectories representing the major classes of CRC (CIN+ and HM).

### refNMF cell-state discovery and deconvolution

Cell states from tumors and normal colonic mucosa were discovered in the VUMC scRNA-seq cohort^9^ using the cNMF Python package^59^. Normal and abnormal epithelial cells plus stromal and immune cells curated from Chen, *et al.* were combined prior to cNMF analysis, and genes were subset to the union of all genes detected in all 40 ST samples^9^. The consensus NMF factorization was performed at an optimal *k* = 30 to yield representative cell-type factors as well as factors that describe cell-state subtypes. Top 500 rank-ordered genes for each cell state are listed in Table S3. We factored the gene loading matrix from scRNA-seq consensus NMF out of the ST expression matrices using the `sklearn.decomposition.NMF` function^103^. This reference NMF (“refNMF”) factorization provided fractional usages of each of the 30 cell-state factors in every ST spot.

### refNMF validation

MxIF stains registered to ST data (Methods: Spatial registration) were averaged within each Visium microwell area for all markers separately. Visium microwells were then blurred using the `squidpy.gr.spatial_neighbors` function with a `radius` value of 1 to capture local spatial neighborhood information of MxIF markers, refNMF cell-state fractions, and gene signature scores^104^. The blurred values were correlated with one another across all Visium microwells from all tumors in the atlas in order to highlight protein markers and cell identity/activity signature scores that confirm refNMF state characterizations (Figure S4E-G).

### Tissue domain detection with MILWRM

We employed refNMF cell states as predictors for a MILWRM model of macro-level consensus tissue domains across all ST slides^63^. We only included states from epithelial and stromal compartments (Figure 4G), excluding immune states to limit predictors to markers of tissue architecture. We used a radius of 1 Visium ST ring for smoothing predictor values, and an *α* of 0.02 for regularization of scaled inertia during *k* optimization.

### Modeling expression dynamics along PPT

We built generalized additive models (GAMs) for genes, gene signatures, and refNMF cell states along global PPT using the `tradeSeq` R package^105^. The pseudotime `(PT)` value given to the model was defined as CNV score for CIN+ tumors and TMB (number of somatic mutations detected per tumor region) for HM tumors (Methods: Global PPT ordering and classification of tumor regions). GAMs were built separately for CIN+ and HM PPT, treating each tumor class as a unique trajectory. We employed the `startVsEndTest` to detect genes, gene signatures, and cell states with differential expression between early and late PPT in both tumor classes, as well as the `diffEndTest` for statistically significant differences in late-PPT expression between the two classes. Resulting significant features from the two tests were curated for heatmap plotting (Figure 4H; Figure 5A; Figure 6A), and `tradeSeq` statistics are provided in Table S4.

### Spatial co-occurrence analysis from ST

We employed the `squidpy` Python package for spatial co-occurrence analysis of IES and infiltrating immune cell states^104^. We first thresholded the signature or cell state of interest to label microwells with “high” expression. Then, we used major CNV clone regions as a reference label to measure co-occurrence with “high” values of the signature or cell state in question. We also employed the stromal (“S”) CNV clone region as a negative control (Figure 5E,K).

### MxIF immune-exclusion analysis

We segmented MxIF images into single cells using the `MIRIAM` segmentation algorithm^76^. Thresholds were manually determined for all markers on a slide-by-slide basis in order to call positive and negative pixels for each marker while avoiding slide-specific illumination or staining batch effects. Within each single cell area, markers were further binarized based on 50 % pixel area or greater, reducing markers to present or absent per single cell segment. Immune cells were then identified by co-expression of the following markers (Figure 5G,I):

- T helper - CD3D and CD4
- T reg - CD3D, CD4, and FOXP3
- T cytotoxic - CD3D and CD8
- Myeloid-CD11B
- Macrophage - CD11B and CD68
- Macrophage M1 - CD11B, CD68, and Lysozyme

ST data for each corresponding slide were transformed into MxIF space using an affine transformation according to `napari`-based spatial registration between MxIF and ST (Methods: Spatial registration). Segmented MxIF data, consisting of cell centroids as x-y pixel coordinates and cell-type IDs (subset to immune cells only, identified with markers above), were then counted for each CNV clone region in the transformed ST data based on centroid pixel coordinates and ST CNV clone masks (Figure 5L-M; Figure S5C).

### TCGA immune exclusion and survival analysis

Normalized expression data (RSEM TPM) and matched clinical information from TCGA COAD and READ samples were accessed and integrated from cBioPortal. Corresponding MMR status (MSS vs. MSI-H) were extracted from GDC Data Portal. Samples with NA values for any of the IES genes were filtered out, yielding 301 total tumors for IES scoring. For stratification by MMR status and survival analysis, these samples were further filtered, removing additional tumors with NA values in any of the metadata columns “Age”, “Gender”, “Race”, or “Disease progressive event and time”. Final sample sizes for survival analysis stratified by MMR status were 244 MSS and 45 MSI-H.

IES scores were calculated as the average TPM expression of *DDR1, TGFBI, PAK4,* and *DPEP1.* Survival analyses were then performed with the `survival` R package (version 3.4-0) (Figure 6B-C; Figure S6A-D).

### Antibody conjugation and immunohistochemical (IHC) imaging

Human CRC TMA slides were cut at 5 μm on positively charged slides. The slides were de-paraffinized in three changes of xylene and hydrated to water in graded alcohol. The slides underwent antigen retrieval using a citrate buffer (Ph 6.0) solution at 105°C in a pressure cooker for 20 minutes followed by a bench cool down for 10 minutes. The slides were washed in distilled water and placed in TBST wash buffer solution for continuation of the staining protocol. Endogenous enzymes were blocked using a 0.03% (H_2_O_2_) peroxidase block solution for 5 minutes, rinsed in wash buffer and incubated for one hour in the primary antibodies (PAK4 Santa Cruz Biotechnology sc-390507 1:75 primary dilution, TGFBI Abcam ab170874 1:300 primary dilution, DDR1 Cell Signaling #5583 primary dilution 1:2000; Key Resources Table). The slides were gently rinsed and a peroxidase labelled polymer was applied utilizing Dako EnVision + System − HRP labeled polymer for a 30 incubation period. After a gentle rinse, the slides were treated with a DAB+ Substrate-Chromogen for 5 minutes to complete the staining protocol. The slides were washed in distilled water, counterstained in Mayer’s Hematoxylin, blued in running tap water, dehydrated in 3 changes of absolute alcohol, and cleared in xylene prior to cover-slipping.

### IHC immune exclusion and survival analysis

From a CRC TMA representing tumors from 163 patients, 149 patients had at least one IHC marker measurement, and 147 of those had clinical data. These tumors were filtered to the subset where DDR1, DPEP1, or TGFBI staining was detected (*n* = 108, 86 MSS and 22 MSI-H). Each stain was graded as 0, 1, 2, or 3 by a pathologist based on increasing expression intensity, and cores were labeled “+” for scores of 2 or 3 and “-” otherwise (Figure 6F). Survival analyses were performed with the `survival` R package (version 3.4-0), comparing “+” to “-” expression for single markers or combinations of markers (Figure 6G; Figure S6E-G).

