## Supplementary material for "Molecular cartography uncovers evolutionary and microenvironmental dynamics in sporadic colorectal tumors": Key Resources Table

| REAGENT or RESOURCE | SOURCE | IDENTIFIER |
| --- | --- | --- |
| Laboratory Reagents | | |
| Mayer’s Hematoxylin | Sigma | MHS16 |
| Alcoholic Eosin | Sigma | HT110116 |
| Bluing Buffer | Dako | CS702 |
| UltraPure Glycerol | Invitrogen | 15514-011 |
| Visium Spatial for FFPE Gene Expression Kit, Human Transcriptome | 10X Genomics | 1000336 |
| DAPI | Sigma-Aldrich | T9284 |
| MUC2 Antibody (F-2) | Santa Cruz | sc-515032 |
| Collagen Antibody (CHP) | 3Helix | R-CHP |
| SNA Antibody (Lectin) | Vector | CL-1305-1 |
| CD11B Antibody (C67F154) | Abcam | ab133357 |
| CD20 Antibody (D-10) | Santa Cruz | sc-393894 |
| PCNA Antibody (PC-10) | Santa Cruz | sc-56 |
| BCATENIN Antibody (12F751) | Vanderbilt Antibody and Protein Resource |  |
| PSTAT3 Antibody (D3A7) | Cell Signaling | 4324S |
| PEGFR Antibody (EP774Y) | Abcam | ab205828 |
| CGA Antibody (C-12) | Santa Cruz | sc-393941 |
| CD4 Antibody (EPR6855) | Abcam | ab133616 |
| COX2 Antibody (D5H5) | Cell Signaling | 13596S |
| CD3D Antibody (EP4426) | Abcam | ab208514 |
| HLAA Antibody (EP1395Y) | Abcam | ab199837 |
| PANCK Antibody (AE1/AE3) | ThermoFisher | 53-9003-82 |
| OLFM4 Antibody (D1E4M) | Cell Signaling | Custom (87240BC) |
| CD8 Antibody (C8/114B) | Biolegend | 13983 |
| ACTININ Antibody (EPR2533(2)) | Abcam | ab198608 |
| CD68 Antibody (KP1) | Santa Cruz | sc-20060 |
| NAKATPASE Antibody (EP1845Y) | Abcam | ab198367 |
| VIMENTIN Antibody (E-5) | Santa Cruz | sc-373717 |
| SOX9 Antibody (EPR14335) | Abcam | ab202516 |
| FOXP3 Antibody (206D) | Biolegend | 320113 |
| LYSOZYME Antibody (E-5) | Santa Cruz | sc-518012 |
| SMA Antibody (1A4) | Santa Cruz | sc-53015 |
| ERBB2 Antibody (MAb414) | Abcam | ab225510 |
| CD45 Antibody (H130) | Biolegend | 304020 |
| ACTG1 Antibody (1-17') | Santa Cruz | sc-65638 |
| MUC5AC Antibody (E309I) | Cell Signaling | Custom (43937BC) |
| CDX2 Antibody (D11D10) | Cell Signaling | Custom (84638BC) |
| DDR1 Antibody (D1G6) | Cell Signaling | 5583 |
| TGFBI Antibody (EPR12078(B)) | Abcam | ab170874 |
| PAK4 Antibody (B-3) | Santa Cruz | sc-390507 |
| Arcturus PicoPure DNA Extraction Kit | Applied Biosystems | KIT0103 |
| truXTRAC FFPE total NA kit | Covaris | 520262 |
| Human Comprehensive Exome Panel | Twist Bioscience | 102033 |
| Human Genome Panel | Twist Bioscience | custom protocol |
| Deposited Data | | |
| Human colon ST | this manuscript | TBD |
| Human colon multiregional WES | this manuscript | TBD |
| Human colon MxIF | this manuscript | TBD |
| Human CRC TMA | this manuscript | TBD |
| Human colon scRNA-seq | Chen *et al.*, 2021 | humantumoratlas.org/explore |
| Human colon bulk WES | Chen *et al.*, 2021 | humantumoratlas.org/explore |
| Human colon MxIF | Chen *et al.*, 2021 | humantumoratlas.org/explore |
| TCGA COAD & READ RNA-seq & clinical data | cBioportal | cbioportal.org |
| TCGA COAD & READ MMR status | NCI GDC Data Portal | portal.gdc.cancer.gov |
| Software and Algorithms | | |
| Space Ranger version 1.3.0 | 10X Genomics | support.10xgenomics.com/spatial-gene-expression/software/downloads |
| Loupe Browser version 6.4.0 | 10X Genomics | 10xgenomics.com/products/loupe-browser/downloads |
| Python version 3.8.10 | Python Software Foundation | python.org |
| R version 4.2.2 | The R Foundation | r-project.org |
| FastQC v0.11.9 | Andrews, 2010 | bioinformatics.babraham.ac.uk/projects/fastqc/ |
| Cutadapt v2.10 | Martin, 2011 | cutadapt.readthedocs.io |
| BWA v0.7.17 | Li & Durbin, 2010 | bio-bwa.sourceforge.net |
| GATK v4.1.8.1 | Van der Auwera & O’Connor, 2020 | gatk.broadinstitute.org |
| Annovar v2019/Dec/05 | Wang, Li, & Hakonarson, 2010 | annovar.openbioinformatics.org |
| Software (Python Packages) | | |
| napari version 0.4.16 | Sofroniew *et al.*, 2022 | napari.org |
| scanpy version 1.9.1 | Wolf, Angerer and Theis, 2018 | github.com/theislab/scanpy |
| matplotlib version 3.0.3 | Hunter, 2007 | matplotlib.org |
| numpy version 1.22.4 | Oliphant, 2006 | numpy.org |
| pandas version 1.4.2 | McKinney *et al.*, 2010 | pandas.pydata.org |
| scipy version 1.8.1 | Oliphant, 2007 | scipy.org |
| seaborn version 0.11.2 | Waskom, *et al.*, 2014 | seaborn.pydata.org |
| scikit-learn version 1.1.1 | Pedregosa *et al.*, 2011 | scikit-learn.org |
| infercnvpy version 0.4.0 | Patel *et al.*, 2014 | github.com/icbi-lab/infercnvpy |
| cnvkit version 0.9.9 | Talevich *et al.*, 2016 | github.com/etal/cnvkit |
| squidpy version 1.2.2 | Palla *et al.*, 2022 | github.com/scverse/squidpy |
| Software (R Packages) | | |
| CytoTRACE version 0.3.3 | Gulati *et al.*, 2020 | cytotrace.stanford.edu |
| maftools version 2.12.0 | Mayakonda *et al.*, 2018 | bioconductor.org/packages/maftools |
| MesKit version 1.6.0 | Liu *et al.*, 2021 | bioconductor.org/packages/MesKit |
| tradeSeq version 1.8.0 | Van den Berge *et al.*, 2020 | bioconductor.org/packages/tradeSeq |
| survival version 3.4-0 | Therneau *et al.*, 2009 | github.com/therneau/survival |
