## Supplemental Figures for "Molecular cartography uncovers evolutionary and microenvironmental dynamics in sporadic colorectal tumors"

### Supplemental information

Key Resources Table

tableS1.csv

**Table S1.** Metadata associated with all spatial CRC atlas samples. WGS and bulk_WES specimens were derived from fresh frozen tissue; all others were derived from FFPE blocks. Empty cells indicate missing or unknown fields.

tableS2.csv

**Table S2.** Gene signatures used for scoring tissue identity and activity in ST and scRNA-seq data. Lists of genes for each signature are utilized by the `scanpy` function `score_genes`. Methods: Gene signature scoring.

tableS3.csv

**Table S3.** Top gene loadings for each of the 30 refNMF cell states discovered in the Chen, *et al.* scRNA-seq cohort. First row contains cell-state ID. 500 genes associated with each cell state are rank-ordered by decreasing spectra score from top to bottom of each column. Methods: refNMF cell-state discovery and deconvolution.

tableS4.csv

**Table S4.** GAM statistics from `tradeSeq` models of top differentially expressed genes, all 30 refNMF cell states, and select gene signatures of interest against PPT for all tumor regions in atlas.

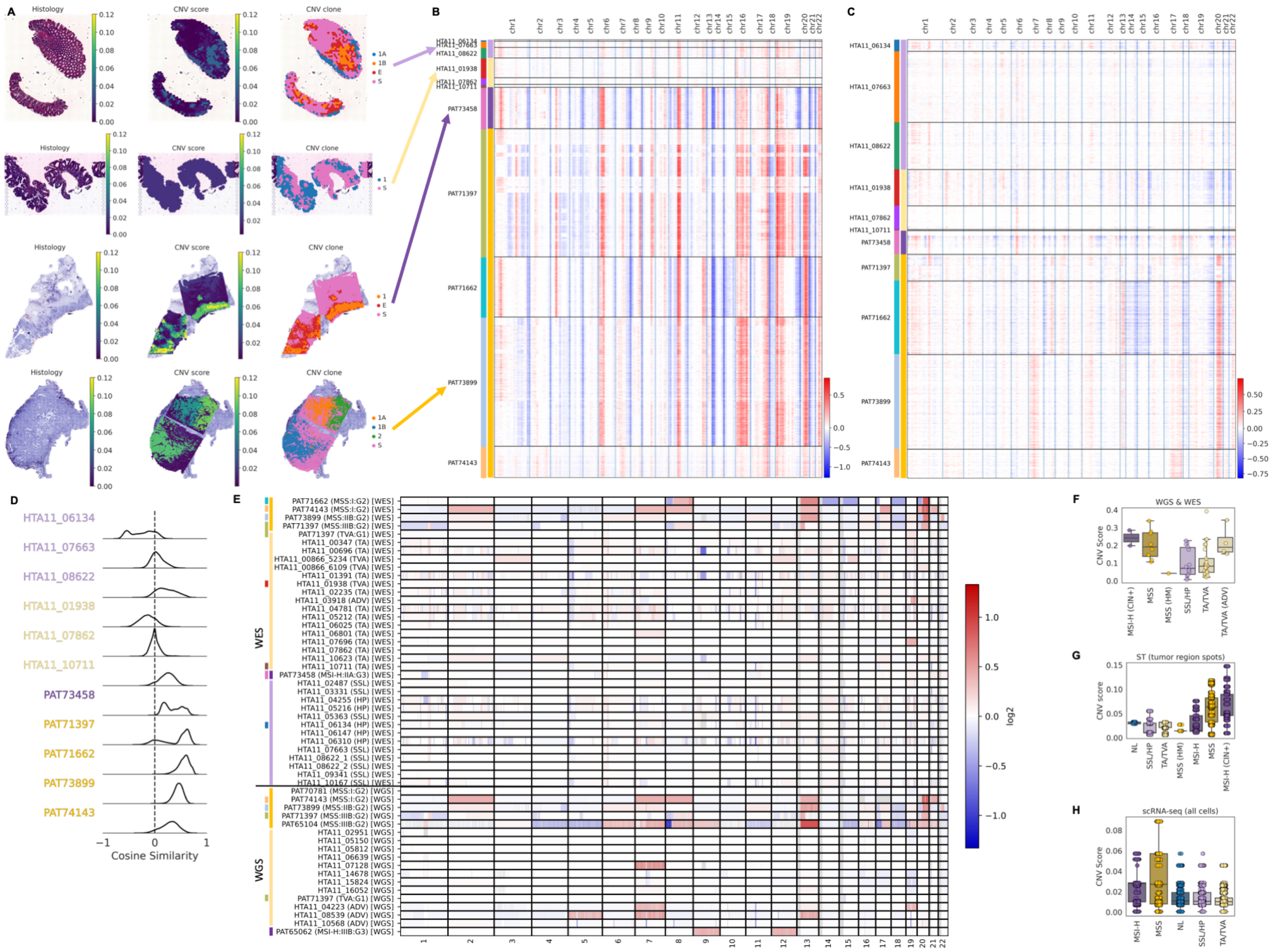

**Figure S2.** CNV inference establishes spatially resolved tumor clones and their phylogenetic relationships.

(A) Representative SSL, TVA, MSI-H, and MSS tumors from the CNV heatmap in B, showing histology alongside CNV score and CNV clone regions.

(B) Heatmap of inferred CNV profiles in the subset of ST samples with matched scRNA-seq. Samples are grouped by patient (left colorbar) and tumor type (right colorbar).

(C) Same as in B for scRNA-seq.

(D) Distribution of cosine similarity scores between the inferred CNV profiles of ST microwells (B) and single cells (C) derived from each patient. Both modalities were subset to major clone regions/clusters prior to comparison to exclude stroma and adjacent normal epithelium (Methods: CNV inference from ST and scRNA-seq).

(E) Heatmap of CNVs detected in WES and WGS of select patients from ST atlas and Chen, *et al.* cohort of pre-cancerous polyps. Samples are colored by patient (left colorbar) and tumor type (right colorbar).

(F) Boxplots of CNV scores for WES and WGS samples in E, grouped by tumor type.

(G) Boxplots of CNV scores for all ST samples, grouped by tumor type.

(H) Boxplots of CNV scores for all scRNA-seq samples, grouped by tumor type.

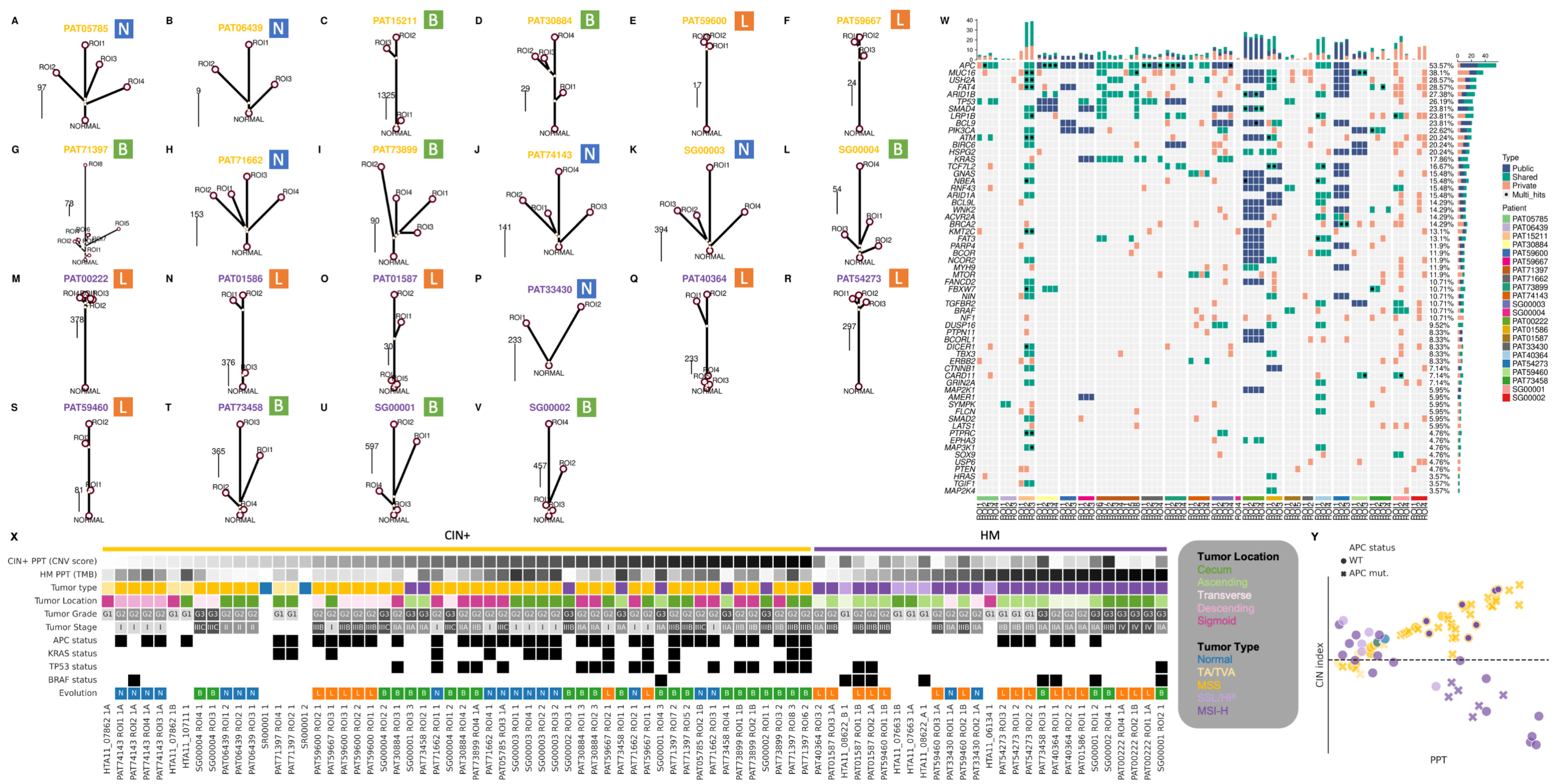

**Figure S3.** Multiregional somatic mutational profiles provide phylogeographical topology.

(A-V) Phylogenetic trees constructed from LCM-WES of 22 CRCs from atlas. Length of branches are proportional to the number of shared or private somatic mutations detected in each region. Patient ID colors represent tumor type (MMR status). Icon next to patient ID indicates inferred mode of evolution based on tree structure ("L" = linear, "B" = branching, "N" = neutral).

(W) Oncoplot of detected driver mutations within the sampled regions of each patient in A-V.

(X) Tumor regions and their clinical and mutational metadata divided by class and ordered left-to-right by corresponding PPT (CNV score for CIN+, TMB for HM).

(Y) CIN index versus PPT for tumor regions from X. Points are colored by tumor class. Dark yellow points with purple centers represent MSI-H/CIN+ tumor regions. Point shape corresponds to regions with detected *APC* mutation.

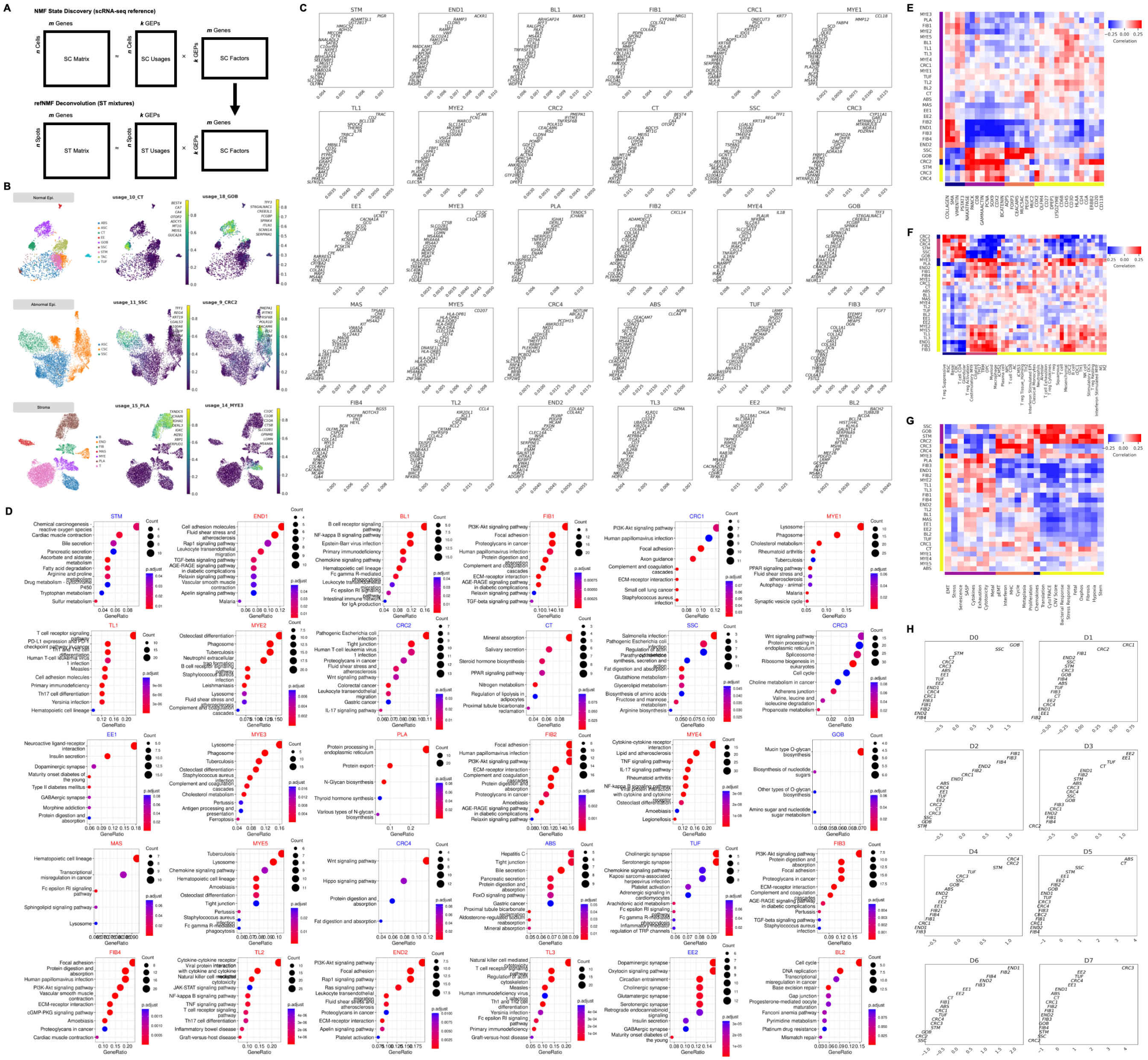

**Figure S4.** Cell-state deconvolution reveals pseudotemporal tissue dynamics.
(A) Diagram of linear algebraic doconvolution of ST expression mixtures using cell states ("factors") discovered by NMF of scRNA-seq. These reference states (SC Factors) are factorized out of the ST matrix to assign fractional scores for all cell states to each microwell (ST Usages).
(B) Example ground-truth NMF factors in Chen, *et al.* scRNA-seq cohort profiling cell states in normal epithelial (top), abnormal/dysplastic epithelial (middle), and stromal (bottom) tissue compartments.
(C) refNMF cell-state gene loadings, showing top 25 genes for each of the 30 consensus states.
(D) KEGG term enrichment for the top 200 genes from each cell state in C. Top 2,000 genes were used for CRC3 to yield associated pathway terms.
(E) Spatial correlation matrix between refNMF cell-state usages (left) and average marker intensity from MxIF data spatially registered to ST from a serial tissue section (bottom).
(F) Same as in E, using cell identity gene signatures.
(G) Same as in E, using cell activity gene signatures.
(H) Tissue domain coefficients for MILWRM model built from all ST data in atlas.

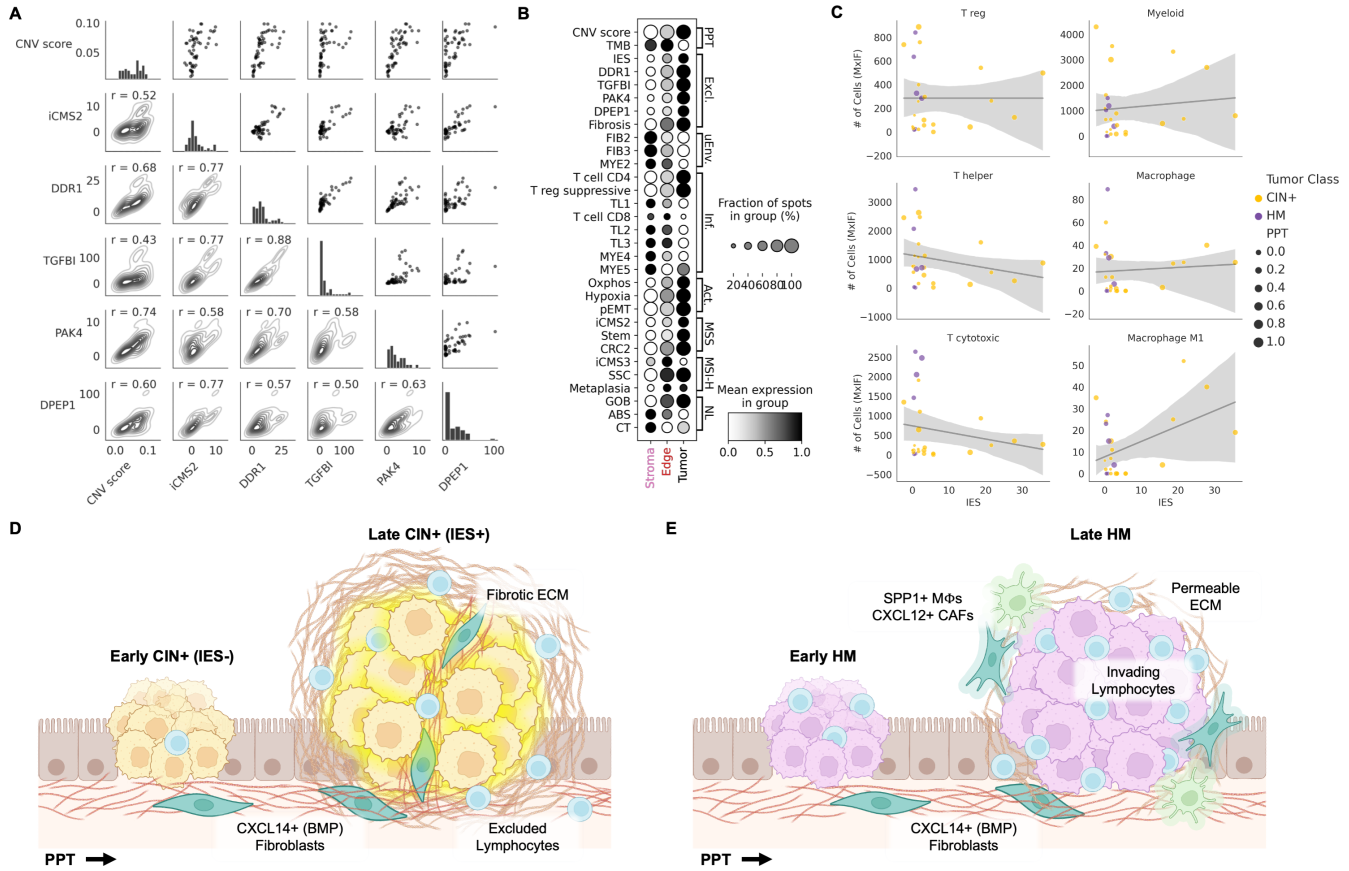


**Figure S5.** Gene expression features of CIN+ CRCs predict immune exclusion.

(A) Pairwise Pearson correlations between IES genes and progression indicators (CNV score and iCMS2) in all CIN+ tumor regions.

(B) Genes, gene signatures, and refNMF cell states grouped into pseudotime indicators ("PPT"), immune exclusion markers ("Excl."), microenvironmental cells ("uEnv."), infiltrating immune cells ("Inf."), tumor activity ("Act."), and epithelial-specific markers of MSS, MSI-H, and normal mucosa summarized across CNV tissue domain for all ST samples.

(C) Number of infiltrating immune cells detected in MxIF plotted against average IES score for all CNV clones in atlas. Points are colored by tumor class and sized according to their respective normalized PPT.

(D) Diagram of early-to-late CIN+ CRC development, highlighting proposed IES mechanism.

(E) Diagram of early-to-late HM CRC.

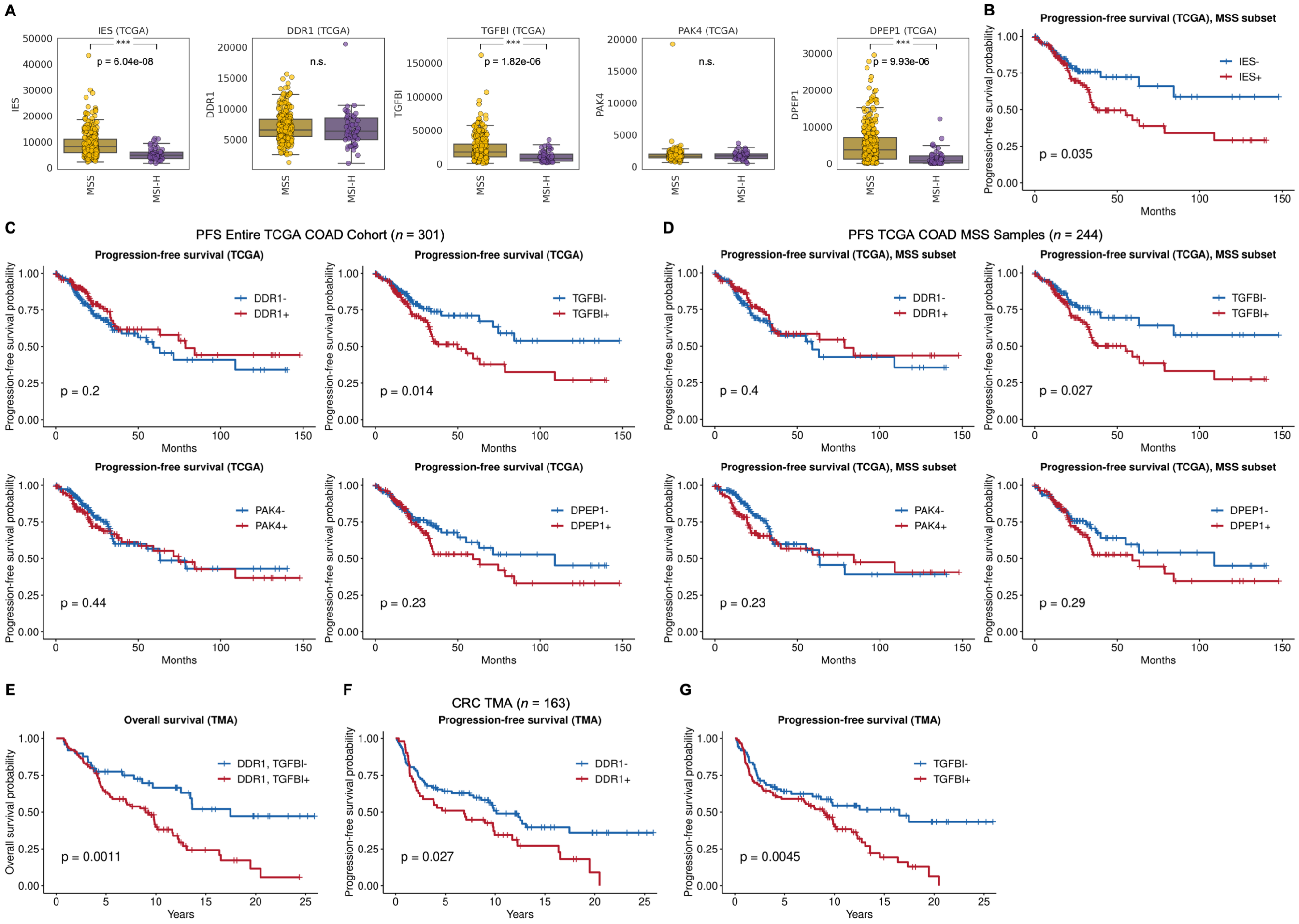


**Figure S6.** IES trends with tumor progression and predicts poor patient outcomes.

(A) Boxplots of IES scores and expression of individual constituent genes in TCGA COAD and READ samples, stratified by MMR status (MSS *n* = 244; MSI-H *n* = 45). Statistics shown represent Student's T-test with Bonferroni correction.

(B) Kaplan-Meier PFS curves for TCGA COAD and READ samples from A with high (+) and low (-) IES scores, subset to MSS tumors only (*n* = 244).

(C) Kaplan-Meier PFS curves for TCGA COAD and READ samples (*n* = 301) with high (+) and low (-) expression of individual constituent genes from IES (*DDR1*, *TGFBI*, *PAK4*, *DPEP1*).

(D) Same as in C, for MSS tumors only (*n* = 244).

(E) Kaplan-Meier PFS curves for CRC TMA cores with high (+) and low (-) DDR1 IHC staining.

(F) Same as in E, for TGFBI.

(G) Kaplan-Meier OS curves for CRC TMA cores with high (+) and low (-) IHC staining of both DDR1 and TGFBI.
